# A platform for multimodal *in vivo* pooled genetic screens reveals regulators of liver function

**DOI:** 10.1101/2024.11.18.624217

**Authors:** Reuben A. Saunders, William E. Allen, Xingjie Pan, Jaspreet Sandhu, Jiaqi Lu, Thomas K. Lau, Karina Smolyar, Zuri A. Sullivan, Catherine Dulac, Jonathan S. Weissman, Xiaowei Zhuang

## Abstract

Organ function requires coordinated activities of thousands of genes in distinct, spatially organized cell types. Understanding the basis of emergent tissue function requires approaches to dissect the genetic control of diverse cellular and tissue phenotypes *in vivo*. Here, we develop paired imaging and sequencing methods to construct large-scale, multi-modal genotype-phenotypes maps in tissue with pooled genetic perturbations. Using imaging, we identify genetic perturbations in individual cells while simultaneously measuring their gene expression and subcellular morphology. Using single-cell sequencing, we measure transcriptomic responses to the same genetic perturbations. We apply this approach to study hundreds of genetic perturbations in the mouse liver. Our study reveals regulators of hepatocyte zonation and liver unfolded protein response, as well as distinct pathways that cause hepatocyte steatosis. Our approach enables new ways of interrogating the genetic basis of complex cellular and organismal physiology and provides crucial training data for emerging machine-learning models of cellular function.

## INTRODUCTION

A central goal of metazoan biology is to understand how coordinated activities of thousands of genes expressed in diverse cell types enables the physiological functions of organs and tissues. Recent large-scale atlases have characterized the molecular composition of a multitude of distinct cell types across the organs ^1–3^; ongoing work is mapping the spatial organization of these cell types across tissues ^4–9^. Although these efforts provide a foundational resource, a major challenge for the field is to causally understand how cell types, cell states, and multicellular networks are produced and regulated by the actions of specific genes and molecular pathways. Addressing this gap in our knowledge will require cellular and spatial functional genomics approaches that enable a systematic and comprehensive understanding of how specific genes control diverse cellular and tissue phenotypes in living animals across physiological states.

Cellular and tissue state manifests across multiple dimensions, ranging from gene expression and post-translational modifications to cellular structures, signaling pathways, and the organization of cells in niches and neighborhoods ^10^. Imaging has historically provided a leading, highly interpretable approach for understanding how genes regulate cellular and tissue state. High resolution imaging of cellular structures in multiple systems has been crucial to innumerable discoveries, ranging from regulators of embryonic development ^11^ to the mechanisms of autophagy ^12^, through the visual identification of phenotypically abnormal cells and organisms. Recent advances in machine learning methods to automate the analysis of complex images has further expanded the power of these approaches ^13^. In parallel, next-generation sequencing has enabled new developments in genome-wide molecular profiling, which provides complementary pictures of cellular state (such as epigenetic, transcriptional, and translational states), in a manner amenable to high-throughput single-cell characterization in tissues.

Coupling these imaging and sequencing tools for deep cell phenotyping to advances in targeted genetic perturbations is markedly expanding our ability to genetically dissect cellular processes at a large scale. In cultured cells, Perturb-seq – the combination of CRISPR screening and single-cell RNA-seq (scRNA-seq) – has enabled the dissection of molecular regulators of core cellular processes such as the unfolded protein response ^14^, hematopoietic development ^15,16^, and alternative polyadenylation ^17^. When applied at the genome-wide scale, this approach has allowed the unsupervised classification of gene function and the principled discovery of cellular responses ^18^. In parallel, advances in highly multiplexed molecular measurements through imaging have enabled new forms of pooled optical screening, using either multiplexed FISH ^19–22^ or *in situ* sequencing ^23–25^ to read out genotypes (i.e., the perturbed gene), including at a genome-wide scale in cultured cells. Such pooled optical screening methods allow the dissection of genes important for phenotypes that require high-resolution imaging to measure.

While *in vitro* studies with cultured cells can dissect many aspects of cellular function, certain aspects of cellular and tissue physiology require study in the context of an intact organism. Technological advances have enabled the generation of mosaic mice harboring a high diversity of perturbation, where individual cells carry distinct perturbations. Alternatively, it is possible to introduce pooled genetic perturbations into animals by transplantation of perturbed cells. The resulting xenograft models have multiple challenges, including clonal bottlenecks, and this approach represents a deviation from the standard animal life cycle. Viral transduction of native tissue has the proven ability to enable genome-scale screens ^26,27^. Using such perturbation delivery approaches, recent pioneering studies in the brain ^28–30^, skin ^31^, and immune system ^15,32^ have used Perturb-seq to profile the effects of perturbing subsets (typically dozens to a few hundred) of disease-relevant genes. However, such sequencing-based approaches do not retain spatial and morphological information such as subcellular structures and tissue organization, which are crucial aspects of cellular and tissue physiology.

A comprehensive understanding that links genes with physiological function will require an integrated approach, bridging multiple modalities of phenotype measurements to obtain a holistic picture of cellular and tissue state in living tissue. Ideally, transcriptionally-defined cellular states would be linked to morphological phenotypes measured through microscopy under different genotypes. By performing a high diversity of genetic perturbations in defined cell types in their native tissues, the function of different genes could be interrogated in their physiological context, under both homeostatic and pathological conditions. Recent imaging-based screening approaches have been expanded to include multi-modal measurements (immunofluorescence and RNA expression) of limited number of genes for cultured cells, or to profile the spatial location of perturbed xenografted cells ^21,33–35^. However, unlike Perturb-seq, these imaging methods have not provided genome-wide transcriptional phenotyping.

Here, we report the development of an approach for performing large-scale pooled genetic screens in native tissue with rich, multimodal phenotypic readouts, as well as its application to map the function of hundreds of genes in the mouse liver under multiple physiological conditions. We are able to study the effects of many genetic perturbations on diverse cellular phenotypes, including transcriptional state, subcellular morphology, and tissue organization, in the liver. Developing this approach requires solving multiple technical challenges towards multimodal, *in vivo* phenotype mapping through both imaging and sequencing. Specifically, we develop new methods for fixed-cell Perturb-seq as well as imaging-based pooled genetic screening in heavily fixed tissue, enabling joint sequencing and imaging analysis of diverse perturbations in the same tissue. For the latter, we apply multiplexed protein and RNA imaging, using *in situ* enzymatic probe amplification followed by multiplexed error-robust FISH (MERFISH) ^36^ to read out both endogenous RNAs and short barcodes for genotyping ^19,21^. Through an integrated analysis of morphological and transcriptional phenotypes, we identify novel regulators of hepatocyte zonation, reveal how proteostatic stress pathway activation can broadly affect the expression of secreted proteins, and show how diverse cellular pathways can produce convergent effects on steatosis. Beyond enabling new ways of interrogating the genetic basis of complex cellular and organismal physiology, this approach will provide crucial training data for emerging machine learning/AI efforts to create predictive models of “virtual” cells^37,38^.

## RESULTS

### Pooled *in vivo* genetic screens through imaging and sequencing

Our goal was to develop an approach that would allow us to profile the effects of a large number of genetic perturbations at subcellular resolution in different cell types in the tissue of a living mouse under defined physiological conditions. For each perturbation, we aimed to measure multiple distinct dimensions of cellular state to capture a more comprehensive picture of gene function. Pioneering work has established the liver as an important system for performing large-scale i*n vivo* pooled genetic screens, due to the effectiveness of viral delivery to hepatocytes in homeostasis ^27^, disease ^39,40^, and regeneration ^41^. These previous studies have focused on cellular survival/proliferation as a phenotype through barcode sequencing. We hypothesized that exploring complex, multimodal phenotypes would allow one to explore a far wider range of gene functions.

To this end, building on recently established ability to deliver genome-wide *in vivo* CRISPR libraries to the mouse liver ^27^, we developed an approach where hepatocytes in a transgenic Cas9 mouse ^42^ were infected in a mosaic with a pool of lentiviruses, each delivering a different CRISPR guide RNAs (Figure 1A). After perfusion with paraformaldehyde to rapidly fix cellular phenotypes, perturbed cells were interrogated using Perturb-Multimodal (Perturb-Multi), a sequencing- and imaging-based assay that captures multiple aspects of cellular state while simultaneously identifying perturbed genes (Figure 1A). From single-cell sequencing, we measured the effects of perturbations on genome-scale transcriptional profiles to obtain rich insights into regulators of cell state in distinct cell type. From imaging, we measured a rich set of phenotypes, including the subcellular morphology and quantity of proteins and mRNAs, as well as the cellular organization of the tissue. By linking cellular properties measured through different modalities via their shared perturbation identity, we then performed integrated multimodal analyses to examine the multifarious effects of the same perturbation on different aspects of cellular function (Figure 1A).

**Figure 1:**
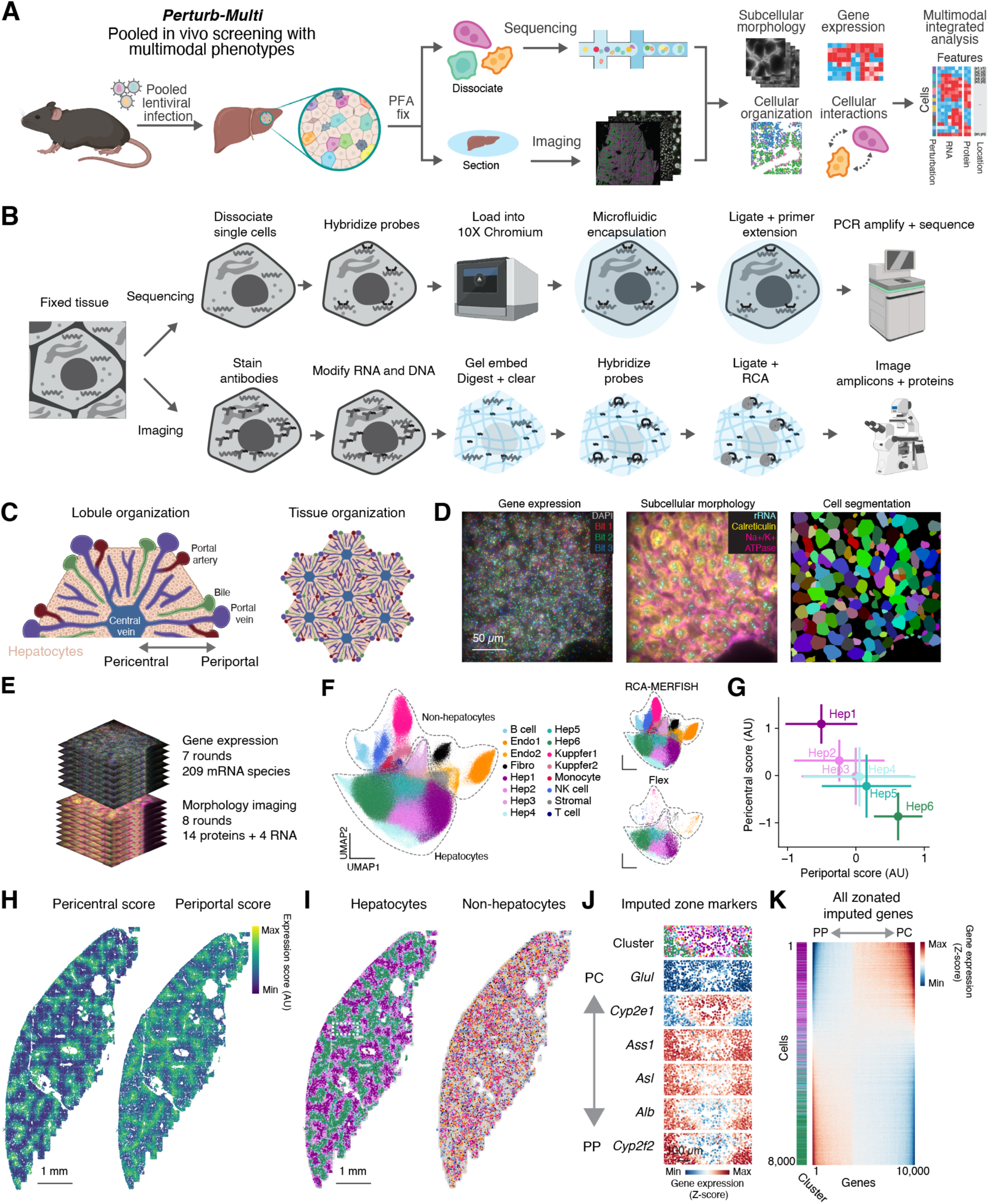
Perturb-Multi: pooled *in vivo* multimodal genetic screens through imaging and sequencing A. Multimodal mapping of genetic and physiological perturbations *in vivo* with Perturb-Multi. Pooled genetic perturbations are introduced into an animal’s liver by lentiviral infection, then animals are fixed with PFA and the liver tissue is subjected either to sequencing- or imaging-based phenotype and genotype readout. Genotype-phenotype relationships on multiple aspects of cellular and tissue function are then interrogated in an integrated analysis. B. Multiple genotype and phenotype readout methods using hybridization-based sequencing and imaging. Top: Sequencing approach. Cells from fixed tissue are dissociated, then subjected to hybridization-based single-cell RNA-seq using 10X Genomics Flex. Bottom: Imaging approach: Tissue sections are stained with pools of oligo conjugated antibodies, then RNA and oligo-antibodies are embedded into an polyacrylamide gel and digested. Probes targeting endogenous RNA and RNA barcodes are hybridized, then ligated and amplified by RCA before readout by MERFISH on an automated microscope. Antibody-associated oligos and abundant RNAs are then detected by sequential rounds of FISH. C. Diagram of liver organization at the lobule and tissue scale, showing major pericentral to periportal axis. Image courtesy of BioRender. D. Multimodal measurement of gene expression (left) and subcellular morphology (center), with machine learning-based segmentation of cells (right). The left panel shows a fluorescence micrograph of an RCA-MERFISH sample stained with DAPI and three fluorescent readout probes for detection of the first three bits. 18 additional bits will be imaged before the mRNA library is decoded. The middle panel shows 3 of the 18 morphological stains that we image in each sample, including two antibodies (anti-Calreticulin, anti-Na+/K+-ATPase) and FISH probe targeting an abundant RNA (pre-rRNA). E. Multimodal readout of RNA and protein through MERFISH (for mRNAs of 209 genes) and sequential rounds of multi-color FISH (for 14 proteins and 4 abundant RNAs). F. Left: Integrated UMAP plot of RCA-MERFISH and Flex single-cell scRNA-seq data, revealing liver cell types identified through unsupervised clustering. Right: same UMAP showing cells from either RCA-MERFISH (upper panel) or Flex scRNA-seq (lower panel). See Methods. G. Periportal vs pericentral gene expression score for hepatocyte subtypes. The periportal and pericentral scores sum the expression of a literature-curated set of periportal or pericentral marker genes, including Cyp2f2 and Hal for periportal score and Glul and Cyp2e1 for pericentral score. H. Spatial distribution of periportal and pericentral scores in hepatocytes. Points represent individual hepatocytes colored by their periportal (left) or pericentral (right) scores. I. Spatial organization of hepatocyte subtypes (left) and non-hepatocyte cell types (right). Points represent individual cells colored by their called cell type or subtype identity. J. Spatial organization of imputed hepatocyte zone markers radially organized around a central vein. Showing zoom from box in (I). The points represent individual cells colored either by their called cell type or subtype identities (top) or by the imputed expression of a gene not directly measured by RCA-MERFISH. The spatial expression patterns are similar to those shown in ^62^. PP: Periportal; PC: Pericentral. K. Genome-wide imputed gene expression of individual hepatocytes (colored by subtype on left). Cells are sorted by periportal gene score, and genes are sorted by correlation with periportal gene score across cells. PP: Periportal; PC: Pericentral.

Measuring phenotypes through both sequencing and imaging required substantial innovation in readout modalities strategies and protocols. For sequencing-based phenotyping, we built on top of the 10X Genomics Flex technology, which uses microfluidic encapsulation of dissociated single cells that have been hybridized with a transcriptome-wide library of split probes against different RNA species for transcriptome-wide detection (Figure 1B, top). Pairs of split probes that hybridize adjacent to each other are then ligated in the droplet and given a cellular barcode for sequencing and expression quantification. We extended this approach to also measure CRISPR-based genetic perturbations through the inclusion of custom probes that directly target the sgRNA.

Creating an imaging approach for genotyping and phenotyping intact tissue sections required extensive technical development. In developing this approach, we had several goals: 1) Create a multimodal protocol that allows for the readout of both RNA and protein from tissue fixed in a manner that is standard for high resolution immunofluorescence, in a single sample preparation on an automated microscope. 2) Amplify RNA FISH signals, both in order to increase the throughput of data collection and to enable the readout of short barcode sequences associated with the sgRNAs. 3) Produce stable samples that could be stored and imaged later, such that many samples could be prepared in parallel and then imaged sequentially.

We explored multiple different approaches, and the approach that we eventually chose and optimized – RCA-MERFISH – builds on several advances over the past several years in methods for spatial transcriptomics and proteomics. For reading out the molecular identity, we used MERFISH ^36,43^, and for signal amplification prior to MERFISH detection, we used rolling circle amplification (RCA) ^44–47^. Specifically, we targeted both endogenous mRNA and barcode RNA (for sgRNA identification) with padlock probes ^48,49^ that can be ligated upon binding to target RNA and amplified *in situ* (Figure 1B, bottom, and Figure S1). Each padlock probe served as an encoding probe and contained multiple readout sequences that together provide a MERFISH code for the target molecule, which can be detected over multiple rounds of hybridization by readout probes complementary to the readout sequences (Figure S1). Because heavy fixation is preferred for immunofluorescence imaging of proteins, to overcome limitations on performing *in situ* enzymatic reactions for RCA in heavily crosslinked tissue, we adopted the hydrogel-based RNA-embedding and tissue clearing approach ^43,45^ and embedding of RCA-amplicons in a polyacrylamide gel ^45,50^. Finally, we combined this approach with multiplexed immunolabeling of proteins with oligo-conjugated antibodies ^51–53^, which were then detected with complementary readout probes using sequential rounds of hybridization. We also detected several abundant cellular RNAs through sequential FISH.

The final protocol consists of a series tissue fixation, embedding, clearing, labeling and imaging steps (Figure 1B). Briefly, after perfusion and cryopreservation, we cut thin sections of tissue, decrosslinked the tissue, and stained it with a pool of oligo-conjugated antibodies targeting different subcellular organelles, membrane, and proteins. After antibody staining, we modified all RNA and DNA in the tissue with an alkylating agent containing an acrylamide moiety (MelphaX) ^54^, and embedded the sample in a thin acrylamide gel, such that these RNA and DNA, including cellular RNAs, sgRNAs with barcodes, and antibody-associated DNA oligonucleotides were covalently linked to the gel. We then digested the proteins and washed away lipids to both clear the tissue and make the remaining RNA and DNA accessible to enzymes and probes. After clearing, we hybridized a library of padlock probes targeting both endogenous RNA and perturbation barcodes. After stringent washing, we then ligated the RNA-hybridized padlock probes and performed rolling circle amplification. The final protocol was the result of extensive optimization of decrosslinking conditions, oligo modifications, additives, and digestion conditions to identify optimal conditions that allow for amplification while being compatible with immunostaining of proteins with oligo-conjugated antibodies in heavily fixed tissue (Figure S2; Methods).

For the RCA-MERFISH detection, each RNA species was targeted by eight padlock probes tiling the transcript and each padlock probe consists of two binding arms that must hybridize adjacent to each other to be ligated, and 4-6 readout sequence for encoding the target RNA (Figures S1C, S1D and S3A). These probes are relatively long (∼140-170 nt) and must be full-length in order to ligate properly. We developed a method to produce these probes from low-cost femtomolar pools of long oligos synthesized in array. After testing multiple different options, we settled on an approach that uses limited cycle PCR and then RCA to amplify the initial array-synthesized oligo pool, followed by Type IIS restriction enzyme digestion to liberate the full length, phosphorylated probes (Figure S3B-E).

### Mapping hepatocyte transcriptional gradients through RCA-MERFISH

As a proof of principle, we applied this imaging approach in combination with scRNA-seq to map the spatial organization of distinct cell types and cell states in the mouse liver. The liver is a classic system for studying cell biology and has enabled discoveries over the past century ranging from the isolation of different organelles to uncovering mechanisms of mTor signaling ^55–58^. For our purposes, the liver provided an excellent test system, as the principal cell type – hepatocytes – have a well-defined molecular composition and spatial organization ^59,60^, and are readily infected with virus for perturbation delivery. At the anatomical level, the liver has long been understood to consist of distinct cellular ‘zones’ radially organized with respect to the central and portal veins (Figure 1C, left) ^61^. Periportal and pericentral hepatocytes have distinct gene expression and correspondingly distinct physiological functions. For example, periportal hepatocytes specialize in oxidative metabolism, gluconeogenesis, and the urea cycle, whereas pericentral hepatocytes specialize in glycolysis and lipogenesis ^61^. This periportal to pericentral distinction is repeated in multiple units around different central veins in a crystalline manner throughout the organ (Figure 1C, right).

After optimization, our multimodal approach allowed us to measure endogenous gene expression while imaging multiple subcellular structures simultaneously. In total, across 7 rounds of 3 color imaging, with a total of 21 bits (Table S1), we measured 209 genes (Table S2) using a Hamming Weight 4 (HW4), Hamming Distance 4 (MHD4) codebook, resolving hundreds of amplicons per cell (Figures 1D, left and 1E). This gene panel was primarily focused on hepatocytes but also included markers for other cell types. In addition to combinatorial RNA imaging with RCA-MERFISH, we sequentially imaged 14 proteins labeling various subcellular organelles, lipid droplets, and signaling pathways and 4 abundant RNA species that exhibited specific patterns of subcellular localization (Figures 1D, middle and 1E). One of our protein targets, the membrane marker Na/K+ ATPase, was used for cell segmentation ^13^ (Figure 1D, right).

Unsupervised clustering revealed a number of hepatocyte subtypes and non-hepatocyte cell types. We classified different cell types as well as subtypes of hepatocytes using an integrated analysis of RCA-MERFISH and scRNA-seq data (Figure 1F). We focused our analysis on hepatocytes. We computed a periportal and pericentral gene expression score based on summing the expression of known marker genes such as *G6pc*, *Aldh1b1*, and *Pck1* ^62^ and found that the hepatocyte clusters fell onto a zonated continuum from Hep1 to Hep6, with Hep1 being the most pericentral and Hep6 being the most periportal (Figure 1G; see Methods). Visualizing these zonation scores and hepatocyte subtypes across the tissue revealed the expected patchwork localization of periportal and pericentral cells (Figures 1H, 1I, and S4), whereas non-hepatocytes had less obvious spatial arrangement (Figure 1I). In addition, we identified one subtype of hepatocytes (Hep3) expressing multiple interferon signaling markers that were localized in small patches (Figure S4; markers include *Isg15*, *Ifit3*, *Stat1*, and *Irf7*). We imputed the expression of the full transcriptome for each cell measured with RCA-MERFISH from the integration with scRNA-seq data and observed the expected pattern of zonated gene expression for imputed genes (Figure 1J, compare with Figure 2 from Ref^62^). Finally, as expected, we found that a large fraction of the transcriptome in hepatocytes varied along the periportal to pericentral axis (Figure 1K).

**Figure 2:**
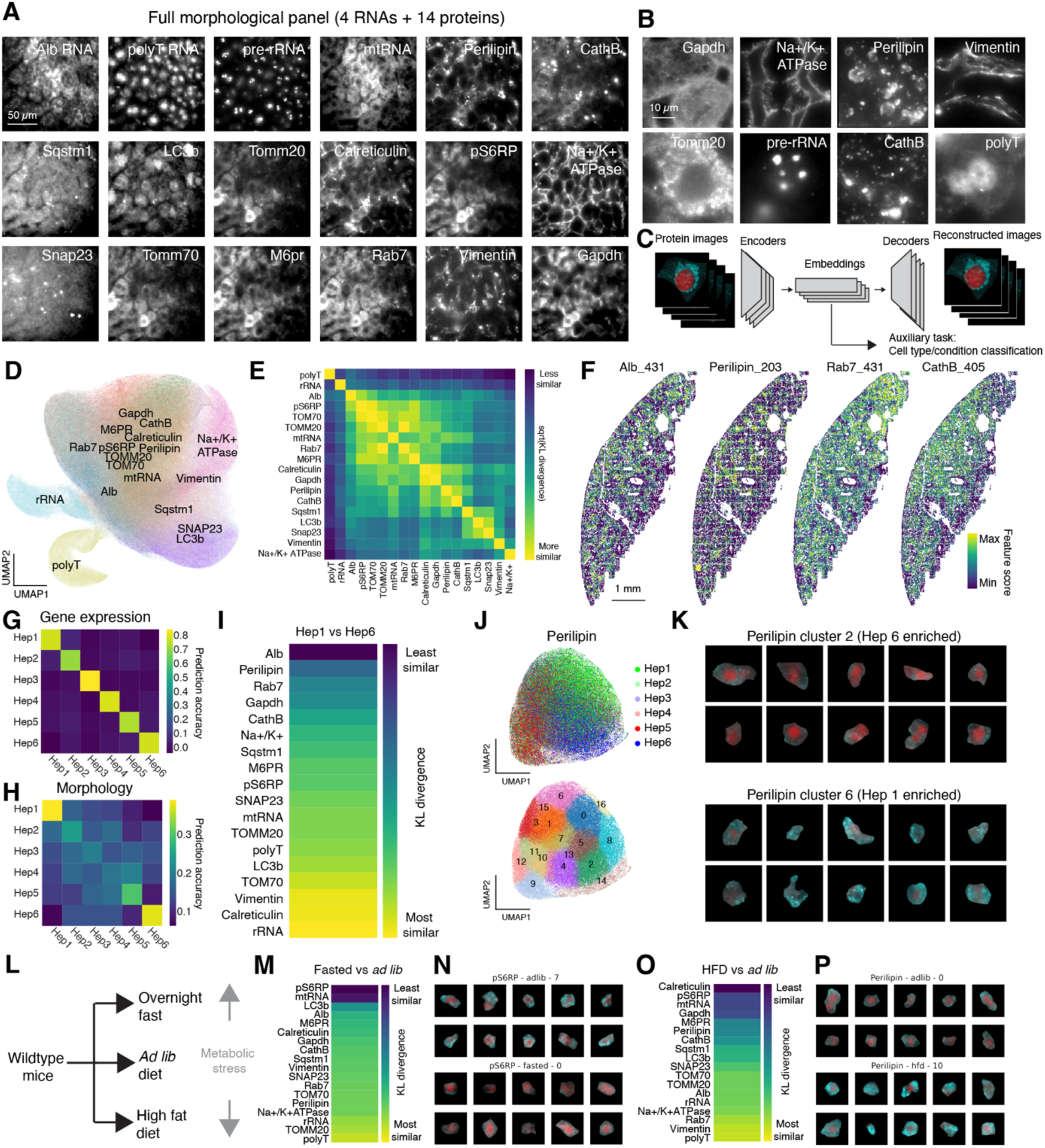
Heterogeneity in subcellular morphology by cell type and state A. Example images of a single field of view imaged with the full subcellular morphology panel, comprising 4 abundant RNA species and 14 proteins. Proteins are labeled with oligo-conjugated antibodies and imaged together with RNAs through sequential rounds of multi-color FISH. B. Zoom-in images of a subset of subcellular morphologies. C. Diagram of deep learning autoencoder to reduce dimensionality and featurize subcellular morphology images. Protein morphologies are reduced from images to a 512 dimensional embedding, using a VQ-VAE model. D. UMAP plot of individual subcellular morphology image embeddings, colored by channel (protein or RNA) identity. The text indicating the morphology channel identity is located at or near (to avoid overlapping text) the centroid of images of the channel in two-dimensional UMAP space. Points represent individual image channels in individual cells, colored by channel identities. E. Similarity of embeddings of subcellular morphological channels, quantified by Kullback Leibler (KL) divergence. Higher KL divergence values indicate a larger difference between the distribution of embeddings. F. Example tissue-scale spatial organizations of single features of morphological embeddings. Each cell in the liver section is colored by its value in the listed feature for a particular protein (for example, the left panel represents *Albumin* mRNA feature 431). G. Confusion matrix representing hepatocyte transcriptional subtype classification accuracy on held out test data, predicted using the transcriptomic data obtained from MERFISH imaging of 209 genes. H. Confusion matrix representing hepatocyte transcriptional subtype classification accuracy on held out test data, predicted using the morphological feature embeddings obtained from imaging the 14 proteins and 4 abundant RNAs. I. Heat map showing mutual information between each morphological channel at hepatocyte subtype identity (Hep1 vs Hep6). Channels with higher KL divergence values have larger differences in the embeddings between Hep1 and Hep6 cells. J. UMAP representation of anti-Perilipin morphological embeddings, colored by hepatocyte subtype identity (top) or embedding Leiden cluster (bottom). K. Unbiased sampling of hepatocytes from Perilipin embedding clusters 2 (enriched for Hep 6 transcriptional subtype) and 6 (enriched for Hep 1 transcriptional subtype). L. Diagram of the diet experiment. M. Heat map showing the capacity of morphological channel embeddings to discriminate between hepatocytes from *ad lib* and fasted mice. Channels with higher KL divergence values have larger differences between the embeddings representing morphology in the *ad lib* and fasted conditions. N. Unbiased sampling of hepatocytes from anti-p-S6 RP embedding cluster 7 (from the *ad lib* condition) and cluster 0 (from the fasted condition). These two clusters have the most significant disparity in enrichment between the conditions and emphasize the difference between conditions. O. Heat map showing the capacity of morphological channel embeddings to discriminate between hepatocytes from *ad lib* and HFD mice. Channels with higher KL divergence values have larger differences between the embeddings representing morphology in the *ad lib* and HFD conditions. P. Unbiased sampling of hepatocytes from anti-perilipin embedding cluster 0 (from the *ad lib* condition) and cluster 10 (from the HFD condition). These two clusters have the most significant disparity in enrichment between the conditions and emphasize the difference between conditions.

### Heterogeneity in subcellular morphology among hepatocytes

To capture diverse aspects of cellular state through high-resolution microscopy, we devised a panel of 18 different protein and RNA targets that covered a variety of subcellular organelles, morphological features such as plasma membrane, and phosphorylated states of proteins. Targeted proteins included both organellar and structural markers such as GAPDH (cytoplasm), Calreticulin (endoplasmic reticulum), Perilipin (lipid droplets), Na+/K+ ATPase (plasma membrane), CathB (lysosome), and Tomm20 (mitochondria), as well as cell state-dependent markers such as the mTOR marker phospo-S6 ribosomal protein (pS6RP) (Table S3). In addition to proteins, we also imaged a few RNA species too abundant to be included in MERFISH imaging, including the highly abundant *Albumin* mRNA, pre-rRNA in the nucleolus, mitochondrial RNA (mtRNA), as well as total polyadenylated RNA (Figures 2A and 2B; Table S4).

To analyze this highly multiplexed cellular imaging data, we built on recent advances in self-supervised learning methods for dimensionality reduction ^63,64^. As part of a data processing pipeline we developed (Figures S5A, S5B), we reduced each image channel (i.e. each protein or abundant RNA target) in each cell from a high-resolution z-stack to a 128x128 pixel matrix and trained a VQ-VAE network ^13^ to reduce the dimensionality of that protein or RNA stain further, to a 512-dimensional vector (Figures 2C, S5C, S5D). The dimensionality reduction was constrained by an auxiliary task that sought to use the low-dimensional embedding to discriminate different transcriptionally-defined cell types or physiological conditions, in order to find an embedding that reflected true biological variation in the sample. We visualized these embeddings with UMAP and found that embeddings representing images of morphologically similar targets were indeed grouped together (Figure 2D); we also observed this through quantification of mutual information between protein/RNA targets (Figure 2E). Further analysis of these embeddings revealed that individual features within the embedding often reflected interpretable patterns (Figure S6; see figure legend and Methods).

Quantifying the variations in subcellular morphology with single-cell resolution at the scale of entire tissues has been a challenging task. This machine-learning-based feature analysis provides a means to overcome this challenge. We took each 512-dimension embedding for a protein/RNA across all cells and visualized individual features from those embeddings. We found that many protein/RNA features exhibited zonal spatial patterns similar to those observed in hepatocyte gene expression (Figure 2F). For some proteins or abundant RNAs, such zonated patterns were expected, such as *Albumin* mRNA and Perilipin ^62^. Others that had zonated profiles to a greater or lesser extent (Rab7, CathB) were less expected. The zonated variation in subcellular morphology of these proteins may reflect differences in hepatic metabolism across different liver zones.

To compare the relative information contained in protein embeddings versus transcriptome in more detail, we attempted to quantify the relative information that was contained in the single-cell transcriptional profiles versus the morphological feature embeddings in the same cells. To do so, we trained a classifier to predict the subtype identity of individual hepatocytes – from Hep1 to Hep6 – on held out cells, using either the 209 genes measured through RCA-MERFISH or the concatenated 18 x 512 dimensional feature embeddings. We found that the transcriptional profiles clearly discriminated all subtypes of hepatocytes (Figure 2G), whereas the morphological feature embeddings only clearly distinguished the most periportal and pericentral subtypes Hep1 and Hep6 (Figure 2H). By contrast, the other subtypes were less clearly distinguished, suggesting that the transcriptionally-defined intermediate subtypes may exhibit a range of subcellular morphologies.

To explore the differences in subcellular morphology between the most periportal and pericentral hepatocyte subtypes, we identified the morphological feature embeddings that best discriminated between Hep1 and Hep6 cells (Figure 2I). *Albumin* mRNA and Perilipin were the most dissimilar between these two hepatocyte subtypes, consistent with the previous understanding of zonated *Albumin* expression ^62^ and lipid storage and metabolism ^61^. We visualized anti-Perilipin embeddings with UMAP and found that embeddings representing cells with different hepatocyte subtype identity were separated (Figure 2J); we conducted unsupervised clustering of the embeddings and found that periportal Hep6 cells were most enriched in anti-Perilipin embedding cluster 2, which has very few lipid droplets, whereas pericentral Hep1 cells were most enriched in anti-Perilipin embedding cluster 6, which has many lipid droplets (Figures 2J and 2K).

To understand the features of liver cellular state that respond to changing physiological conditions and metabolic stress, we overnight-fasted mice or fed them a high-fat diet (HFD) for a month and fixed tissue samples for analysis by scRNA-seq and imaging (Figure 2L). These physiological perturbations caused large changes in gene expression, especially in hepatocytes (Figure S7A). We embedded the morphological images with our VQ-VAE model and looked for image channels whose embeddings best discriminated between these two physiological conditions. Phospho-S6 ribosomal protein (pS6RP) staining separated hepatocytes between the fasted and *ad libitum* (*ad lib*) samples best, consistent with S6 ribosomal protein phosphorylation by mTOR in response to nutritional status (Figures 2M, 2N). Perilipin differed in cells from the *ad lib* and HFD samples (Figures 2O, 2P). We note that stains such as anti-calreticulin and anti-pS6RP were also quite different between the *ad lib* and HFD samples. This may in part reflect negative space caused by the exclusion of these stains by large lipid droplets, in addition to potential changes in the abundance of these targets caused by the altered dietary conditions (Figures S7B, S7C).

In total, these experiments demonstrated that our multiplexed protein and RNA imaging, combined with deep learning analysis, was able to resolve significant heterogeneity in subcellular morphology, which could be further linked to distinctions in transcriptionally-defined cell types and the animal’s physiological state. These experiments create the first integrated transcriptional and subcellular morphological map of the liver, while validating the ability of protein and RNA readouts to provide complementary information on liver anatomy and physiology.

### Large-scale *in vivo* multimodal screening in CRISPR mosaic livers

Having established an integrated sequencing and imaging methodology for studying the molecular and cellular organization of tissue at multiple scales, we combined this approach with large-scale pooled *in vivo* CRISPR screening to map the effect of genetic knockout perturbations on genome-wide transcriptional state as well as the intensity and morphology of the imaged proteins and RNAs. We targeted 202 genes (Table S6), each with two independent sgRNAs, and included 50 negative control sgRNAs. The genes chosen for perturbation were curated from the literature and were involved in diverse aspects of hepatocyte and liver physiology, including metabolism, intercellular and intracellular signaling, transcription and translation, and protein trafficking and secretion. We also targeted genes selected for high or specific expression in hepatocytes, including genes with uncertain functions in hepatocytes.

We introduced barcoded genetic perturbations that could be read out through either imaging or sequencing into the liver through lentiviral delivery. Building on the CROP-seq lentiviral vector design ^65^ to mitigate the effects of recombination during lentivirus production, we engineered a vector that expresses a fluorescent protein (mTurquoise) as well as a 185-mer barcode associated with a sgRNA for imaging from a hepatocyte-specific Pol II promoter while simultaneously expressing the sgRNA from a Pol III promoter (Figure 3A). Each sgRNA and its associated 185-mer barcode were synthesized on a single DNA fragment and then cloned into the vector in a pool. An intervening region composed of one of three alternative sgRNA constant regions was introduced between the barcode and sgRNA to reduce the effects of barcode swapping through recombination of the constant intervening sequence ^14,66^.

**Figure 3:**
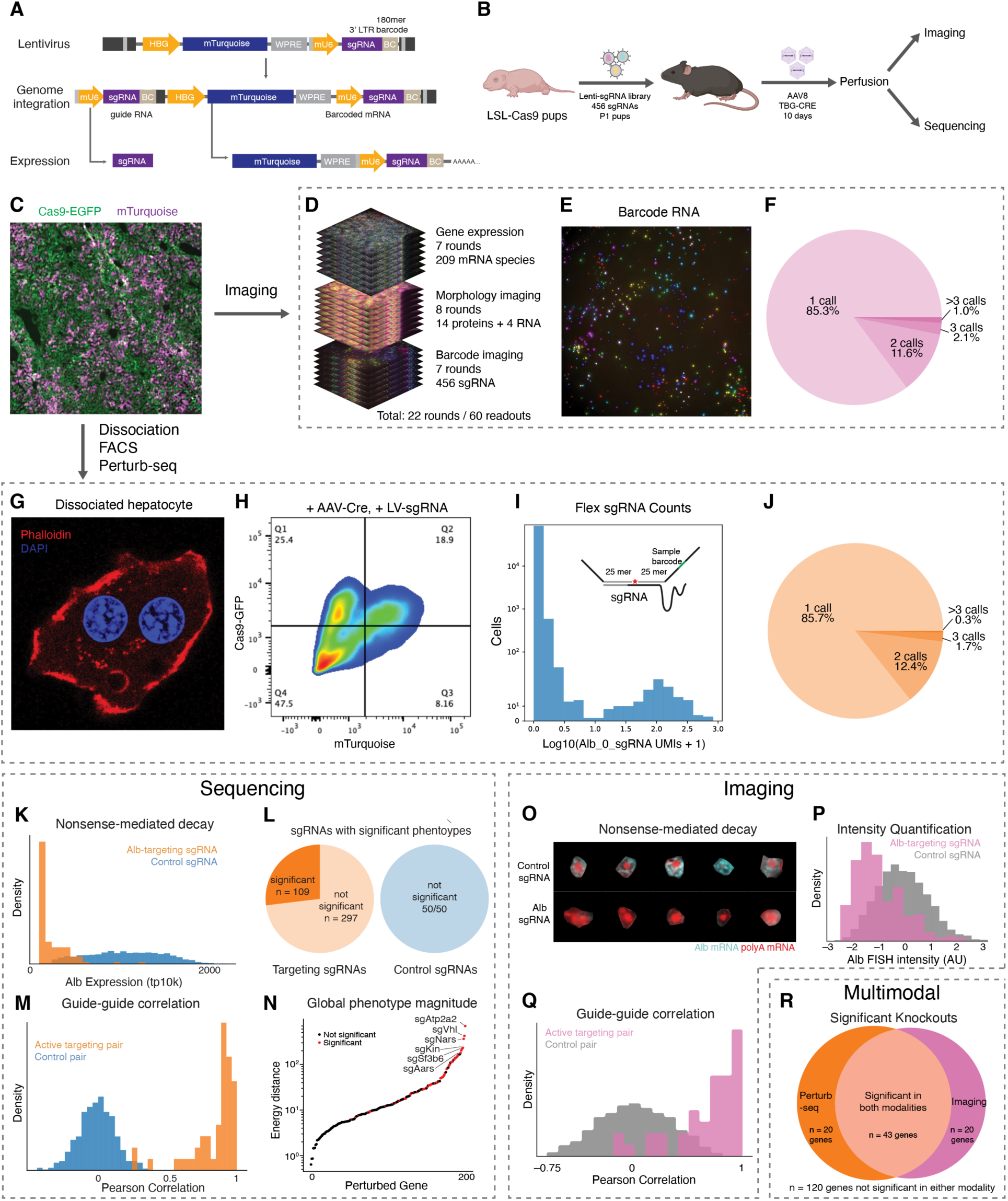
Large-scale, multimodal *in vivo* screening in CRISPR mosaic livers A. Schematic of lentiviral vector used for dual-mode mosaic screens. The vector was derived from a CROP-seq vector and includes an mU6 promoter driving sgRNA expression and a hepatocyte promoter driving expression of an mTurquoise transcript that contains a perturbation-specific barcode sequence in the 3’ UTR. B. Diagram of CRISPR perturbation experiment. LSL-Cas9 pups are injected with an sgRNA perturbation library. Cas9 is activated in young adults through the administration of AAV8 TBG-CRE and livers are harvested with PFA perfusion and then analyzed with RCA-MERFISH or fixed cell Perturb-seq. C. Fluorescence micrograph of PFA-perfused, lentivirus- and AAV-transduced liver tissue. The micrograph shows cells expressing Cas9-EGFP (green) and sgRNA-mTurquoise (purple). D. Multimodal readout of RNA, protein, and perturbation barcode through sequential hybridization and imaging. 209 endogenous RNAs and 456 perturbation barcodes are imaged with two back-to-back RCA-MERFISH runs on the same instrument, alongside the imaging of 14 proteins and 4 abundant RNAs with sequential rounds of multicolor-FISH. E. Representative fluorescence micrograph of the first three bits (out of 21 bits total) of RCA-MERFISH imaging of perturbations. F. Pie chart representing the number of sgRNA calls in each cell in the imaging experiment. 85.3% and 14.7% of imaged cells with at least one called perturbation barcode have exactly one and two called barcodes, respectively. In this study, we only analyzed cells with a single called perturbation. G. Fluorescence micrograph of a representative hepatocyte recovered after the dissociation of PFA-perfused liver. The colors represent DAPI and phalloidin. H. Representative flow cytometry data from dissociated PFA-perfused, lentivirus- and AAV-transduced liver tissue. Cells are sorted for mTurquoise+ and GFP+ signals to enrich for a population that received an sgRNA and has active Cas9. I. Histogram representing counts data from custom hybridization probes targeting one of the two *Albumin* sgRNAs, Alb_0. The y axis is semi-log. The inset shows a schematic of direct split probe hybridization to the variable and constant portions of an sgRNA. J. Pie chart representing the number of sgRNA calls in each cell in the Perturb-seq dataset. 85.7% and 14.3% of imaged cells with at least one called perturbation barcode over the threshold have exactly one and two called barcodes, respectively. K. Histogram comparing *Albumin* expression in called cells with a control sgRNA and called cells with *Albumin*-targeting sgRNAs, from the Perturb-seq dataset. The x axis represents the tp10k expression of *Albumin*. L. Pie charts showing the number of targeting (left) and non-targeting (right) sgRNAs that caused a significant transcriptional phenotype, as measured by a Holm-Sidak-corrected energy distance permutation test (p < 0.05), in the Perturb-seq dataset. 109/406 targeting sgRNAs (27%) and 0/50 of non-targeting sgRNAs (0%) have significant transcriptional phenotypes. M. Histogram comparing the Pearson correlation of pseudo-bulk transcriptional changes associated with pairs of active targeting sgRNAs targeting the same gene, to pairs of control sgRNAs, in the Perturb-seq dataset. N. Ranking knockouts according to the energy distance vs control cells, in the Perturb-seq dataset. The energy distance is calculated using the top 20 PCs of Z-normalized gene expression. O. Unbiased sampling of cells with control sgRNAs and sgRNAs targeting *Albumin*. The fluorescence micrographs show *Albumin* mRNA and polyA FISH channels. P. Histogram comparing *Albumin* mRNA intensity in called cells with a control sgRNA and called cells with *Albumin*-targeting sgRNAs, from the imaging dataset. The x axis represents the intensity of *Albumin* mRNA FISH across each individual cell, relative to the average intensity of *Albumin* FISH in cells with control sgRNAs. Q. Histogram comparing the Pearson correlation coefficients of pseudo-bulk transcriptional changes associated with pairs of active targeting sgRNAs targeting the same gene, to pairs of control sgRNAs, in the imaging dataset. R. Venn diagram of genes with significant knockout phenotypes in imaging and sequencing. Phenotype significance in the imaging and sequencing assays are measured by a Holm-Sidak-corrected energy distance permutation tests (p < 0.05). There is significant overlap in the two sets of genes (hypergeometric p < 10^-13^).

We then used this vector to introduce a mosaic of genetic perturbations into the liver of Cas9 transgenic mice ^42^. We produced high-titer pools of CROP-seq lentivirus and introduced the virus systemically into transgenic Cre-inducible Cas9 mice at postnatal day 1 (P1) through injection into the temporal vein (Figure 3B). We targeted a relatively low MOI (10-30%) such that most infected cells had a single perturbation. We then allowed the mice to grow to adulthood (>P30) and induced Cas9 expression in hepatocytes with systemically delivered AAV containing the Cre recombinase driven by a hepatocyte-specific TBG promoter. After waiting 10 days to allow Cas9 to express and perturb the targeted gene, the mice were perfused and the fixed livers were cryopreserved for analysis. Lentivirus introduced at P1 expressed mTurquoise in mosaic distribution throughout the liver of adult mice, whereas EGFP expressed with Cas9 was uniformly expressed in presumptive hepatocytes following Cre recombination (Figure 3C). We then mapped the effects of these perturbations on hepatocytes using a combination of imaging and sequencing.

For imaging-based phenotype and genotype measurement, we extended the RNA and morphological imaging approach described above with the inclusion of padlock probes targeting the perturbation-specific barcode sequences, which are orthogonal to the mouse genome. These perturbation-specific probes carry the readout sequences that encode the perturbations (i.e., in the present studies the sgRNA, but this could be extended to other genetically-encoded perturbations) with a HW6, HD4 MERFISH code, which were imaged by MERFISH alongside the endogenous mRNA imaging and sequential imaging of proteins and abundant RNAs on the same automated microscope (Figures 3D and 3E).

As expected, sgRNA-associated barcodes were expressed only within a subset of cells. After establishing a count threshold to confidently identify cells with each perturbation, we found that most cells did not contain any perturbation and that most perturbed cells had only a single perturbation, consistent with the low infection MOI and approximately Poisson-distributed transduction of the lentivirus-accessible cells in the liver (Figure 3F). We observed relatively homogeneous infection throughout the tissue and within lobules. Altogether, we analyzed ∼79,000 cells perturbed with single sgRNAs.

In parallel, to measure genome-wide transcriptional responses to the genetic perturbations, we developed a fixed-cell Perturb-seq method. In addition to compatibility with imaging, we reasoned that a fixed cell protocol could overcome many of the challenges that have limited the scale and practicality of *in vivo* Perturb-seq studies, such as sample degradation during tissue dissociation and FACS enrichment ^67,68^, especially when processing many tissues in parallel or recovering millions of cells.

Dissociation of fixed tissue from the same livers that we imaged yielded intact cells with morphologies similar to those observed in tissue (Figure 3G). Despite heavy fixation, the cells could be efficiently dissociated with high yield, obviating the need for nuclear RNA sequencing ^69^ or complex enzymatic dissociation approaches ^70^. We used FACS to enrich for sgRNA-transduced (mTurquoise+) and Cas9-active (GFP+) cells (Figure 3H) and then performed fixed-cell single-cell RNA-seq using the hybridization-based 10X Flex approach. To determine the identity of expressed sgRNA alongside the mRNA transcriptome, we spike in a pool of custom probes that directly targeted the protospacers of the sgRNA library (right hand side probe) and the constant region of the sgRNA (left hand side probe). In most cells, one or two sgRNA probes had much higher UMI counts than other sgRNAs, most of which gave zero UMI counts (Figure 3I). We used this distribution to establish a threshold and confidently identify the sgRNAs present in each cell. Most perturbed cells had a single sgRNA above threshold, with only a small fraction of cells harboring two or more distinct sgRNAs (Figure 3J), again consistent with the low MOI. Altogether, we analyzed ∼55,000 cells perturbed with single sgRNAs.

### Validation of perturbation calling

We confirmed the accuracy of our sgRNA calling using Perturb-seq and RCA-MERFISH by inspecting for depletion of sgRNA targets. Fortuitously, sgRNAs targeting *Albumin* (*Alb*), an abundant secreted protein whose mRNA provides ∼10% of the mRNA UMIs in hepatocytes, caused strong depletion of *Albumin* mRNA relative to control sgRNAs, presumably due nonsense-mediated decay (NMD) of targeted mRNAs. With Perturb-seq, we quantified *Albumin* mRNA depletion directly (Figure 3K) and observed minimal overlap between the distributions of *Albumin* expression in cells expressing sgRNAs against *Albumin* and those expressing control sgRNA, indicating a low false positive rate of sgRNA calling. We also conducted a fixed-cell Perturb-seq experiment in cultured CRISPRi K562 cells, which allowed us to measure on-target knockdown without relying on NMD (median 87% knockdown across targets, Figure S8), confirming the accuracy and specificity of sgRNA calling with hybridization-based scRNA-seq.

Because probes against *Albumin* mRNA were also included in our imaging-based screens, we quantified raw FISH signal for *Albumin* mRNA and observed lower intensity in cells expressing sgRNAs against *Albumin* as compared to cells expressing control sgRNA (Figures 3O and 3P). Likewise, our immunofluorescence imaging showed that anti-Gapdh signals were substantially lower in cells expressing sgRNAs against *Gapdh* than in cells expressing control sgRNA (Figures S9A and S9B). These results suggest that perturbation readout by RCA-MERFISH also had a low false positive rate.

Next, we explored the broader phenotypic consequences of each genetic perturbation measured by Perturb-seq and imaging. As a first-pass analysis of the imaging data, we quantified the mean intensity of each 14 proteins or 4 abundant RNA targets in each cell.

We used an energy-distance permutation test with stringent multiple testing correction ^71^ to determine whether perturbations caused a difference in the distribution of (1) global transcriptional states measured by Perturb-seq and (2) overall morphological states measured by imaging, relative to cells with control sgRNAs. Such global analyses are crucial, as experiments perturbing hundreds of genes and measuring tens of thousands of phenotypes across a multitude of cells can be prone to statistical false discovery ^71^. We note that, although we designed our library to be enriched for targets with functions in liver biology, we do not have ground-truth knowledge of which sgRNAs can cause phenotype changes in our assays. In the Perturb-seq experiment, 109/406 targeting sgRNAs had a statistically significant impact on the distribution of transcriptional states (measured as the first 20 PCs of mRNA expression) (Figure 3L); at that significance threshold (multiple-testing corrected p value < 0.05), 0/50 control sgRNAs had a significant transcriptional impact. In the imaging experiment, 84/406 targeting sgRNAs and 3/50 control sgRNAs had a significant impact on the distribution of morphological state (Figure S9C). Furthermore, we compared pairs of sgRNAs targeting the same gene. We generated pseudo-bulk sgRNA-phenotype maps by groups cells with identical sgRNAs and found that pairs of sgRNAs targeting the same gene caused both highly correlated transcriptional phenotypes and highly correlated morphological phenotypes, relative to random pairs of non-targeting sgRNAs (Figures 3M and 3Q). Together, these results support the accuracy of our sgRNA calling and specificity of the perturbations caused by these sgRNAs.

We inspected the perturbations that had the largest impact on the distribution of global phenotypic states, reflected in energy distance from cells with control sgRNAs (Figures 3N and 3R). In the Perturb-seq experiment, many genes with functions essential for cellular growth and survival, such as *Atp2a2*, *Aars*, and *Sf3b6* caused large impacts to global transcriptional state upon acute knockout (Figure 3N). The knockout of signaling genes such as *Vhl* also had large transcriptional impacts (Figure 3N). Many of the same genes also had large impacts on the global immunofluorescence state (Figure 3R; hypergeometric p < 10^-13^). Overall, we observed a positive correlation between the magnitude of the knockout phenotypes in the Perturb-seq and imaging assays (Figure S9D, Pearson’s R = 0.5). We note that since our Perturb-seq and imaging experiments measured different phenotypes—genome-wide transcription in the former and 14 proteins and 4 abundant RNAs in the latter—we do not necessarily anticipate a high correlation between these two measurements.

Next, we identified the differentially expressed genes associated with each sgRNA. Although most sgRNAs caused differential expression in a relatively small number genes, comparable to the effects of control sgRNA, some targeting sgRNAs caused hundreds to thousands of genes to be differentially expressed in the Perturb-seq measurements (Figure S9E), consistent with our earlier observation that ∼25% of targeting sgRNAs had a significant impact on the transcriptional state (Figure 3L). We also identified the protein and abundant RNA channels that were significantly impacted by each perturbation in the imaging measurements. Most active targeting perturbations impacted a relatively small subset of imaged targets, but some perturbations such as the essential nuclear genes *Sf3b6*, *Sbno1*, *Polr1a*, and *Kin* had a significant impact on many channels (Figure S9F). These results presumably reflect the pleiotropic consequences of disrupted transcription and RNA processing.

### Multimodal *in vivo* screening with strong phenotypes

Our imaging and sequencing experiments resulted in >100,000 total perturbed cells with either 18-plex imaging phenotypes (subcellular morphology and cell locations in tissue), or genome-wide transcriptional phenotypes. We visualized the spatial organization of perturbed cells in tissue and observed small clusters of cells sharing the same perturbation, distributed throughout the various zones that form liver tissue (Figure 4A); these clusters presumably reflect local proliferation of single infected hepatocytes during postnatal growth after lentiviral transduction. We visualized the Perturb-seq transcriptional phenotypes with dimensionality reduction. For example, a UMAP projection of transcriptional phenotypes clearly separated cells with control sgRNAs from cells with sgRNAs targeting the core hepatocyte transcription factor Hnf4a (Figure 4B). We noted that *Hnf4a* knockout was associated with lower expression of Apolipoprotein AI, consistent with the role of *Hnf4a* in promoting the expression of abundant secreted factors in hepatocytes^72^.

**Figure 4:**
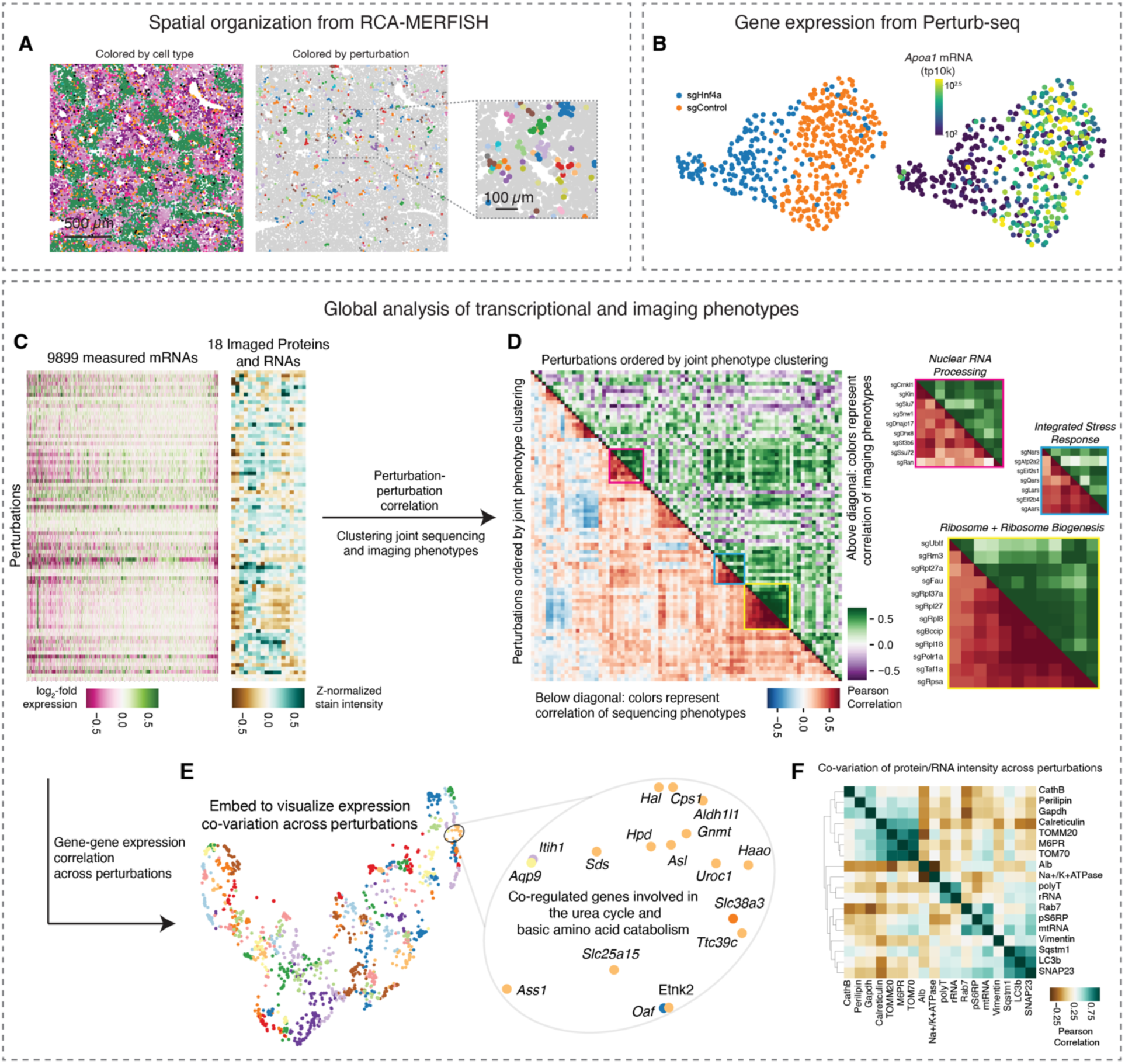
Multimodal *in vivo* screening with strong phenotypes A. An example exploration of a spatial phenotype in the imaging dataset. The cells are colored by cell type (left; the colors are the same as in Fig. 1F) or by sgRNA barcode identity (right and zoom; cells with no called sgRNA are light gray). B. An example exploration of a transcriptional phenotype in the Perturb-seq dataset. A UMAP representing individual cells is generated from the single cell transcriptomes of cells with sgRNAs targeting *Hnf4a* and from a random sub-sampling of cells with control sgRNAs. The UMAP is colored by sgRNA identity (left) or by *Apoa1* expression (right). C. Heat map representation of pseudobulk transcriptional changes (measured by sequencing, left) and staining protein and RNA level changes (measured by imaging, right) associated with each sgRNA, relative to cells with control sgRNAs. The colors represent perturbed gene-level log2-fold RNA expression changes in cells with perturbed genes versus negative control cells (transcriptional sequencing data), or Z-normalized protein/RNA expression change in cells with perturbed genes relative to negative control cells (staining intensity imaging data), and are clipped for visual emphasis. D. Heat map of the perturbation-perturbation correlation of RNA and protein changes associated with active sgRNAs, with zooms. The colors represent Pearson correlation of perturbed gene-level pseudobulk phenotypes in the sequencing dataset (below diagonal) or the imaging dataset (above diagonal). The genetic perturbations are ordered by single linkage hierarchical clustering of joint phenotype vectors that represent both transcriptional changes measured by sequencing and protein and RNA changes measured by imaging. E. Minimal distortion embedding where each dot represents an mRNA expressed in hepatocytes. mRNAs that are co-varying in expression across the genetic perturbations are placed in proximity. F. Heat map of the correlation between the expression levels of indicated proteins/RNAs across perturbations, in the imaging dataset. The proteins/RNAs are ordered by single linkage hierarchical clustering of the correlation matrix.

We performed unbiased clustering of the multimodal genotype-phenotype map to identify perturbations that cause similar multimodal phenotypes measured by Perturb-seq and multiplexed imaging (Figures 4C and 4D). The perturbation cluster map derived from joint analysis of both modalities recovered many interpretable blocks of perturbed gene function, including a large block of ribosomal proteins and ribosome biogenesis genes, a block of nuclear mRNA processing genes, and a block of genes whose disruption activates the integrated stress response ^18^ (Figure 4D). Broadly, the sequencing and imaging data identified related but complementary perturbation-perturbation relationships (Pearson’s *R* = 0.42 for correlation-of-correlations).

We identified mRNA and protein phenotypes that co-varied across perturbations. We visualized the mRNA co-expression data from sequencing by constructing a minimum distortion embedding that placed genes with correlated expression nearby (Figure 4E; Methods). We identified clusters of co-regulated genes with similar biological functions, such as a group of genes involved in the urea cycle and the transport and metabolism of basic amino acids. We also identified imaging stains whose intensity co-varied across perturbations (Figure 4F), such as the mitochondria proteins Tomm20 and Tomm70, the lysosomal protein mannose 6-phosphate receptors (M6PRs), and the ER chaperone calreticulin.

### Identifying genetic regulators of liver physiology

We conducted targeted analyses to discover key regulators of diverse aspects of *in vivo* liver physiology. With the Perturb-seq data, we identified the drivers of gene expression modules with crucial functions in hepatocytes. For example, we ranked genetic perturbations by their impact on a core program of lipid biosynthesis genes defined by a previous large-scale Perturb-seq study ^18^ (Figure 5A; see methods). Knockout of the lipid biosynthesis inhibitor *Insig1* had the strongest positive effect on lipid biosynthesis gene expression; knockout of the Ldl receptor (*Ldlr*) and *Srebf1* also had strong positive impacts, possibly due to compensatory expression.

**Figure 5:**
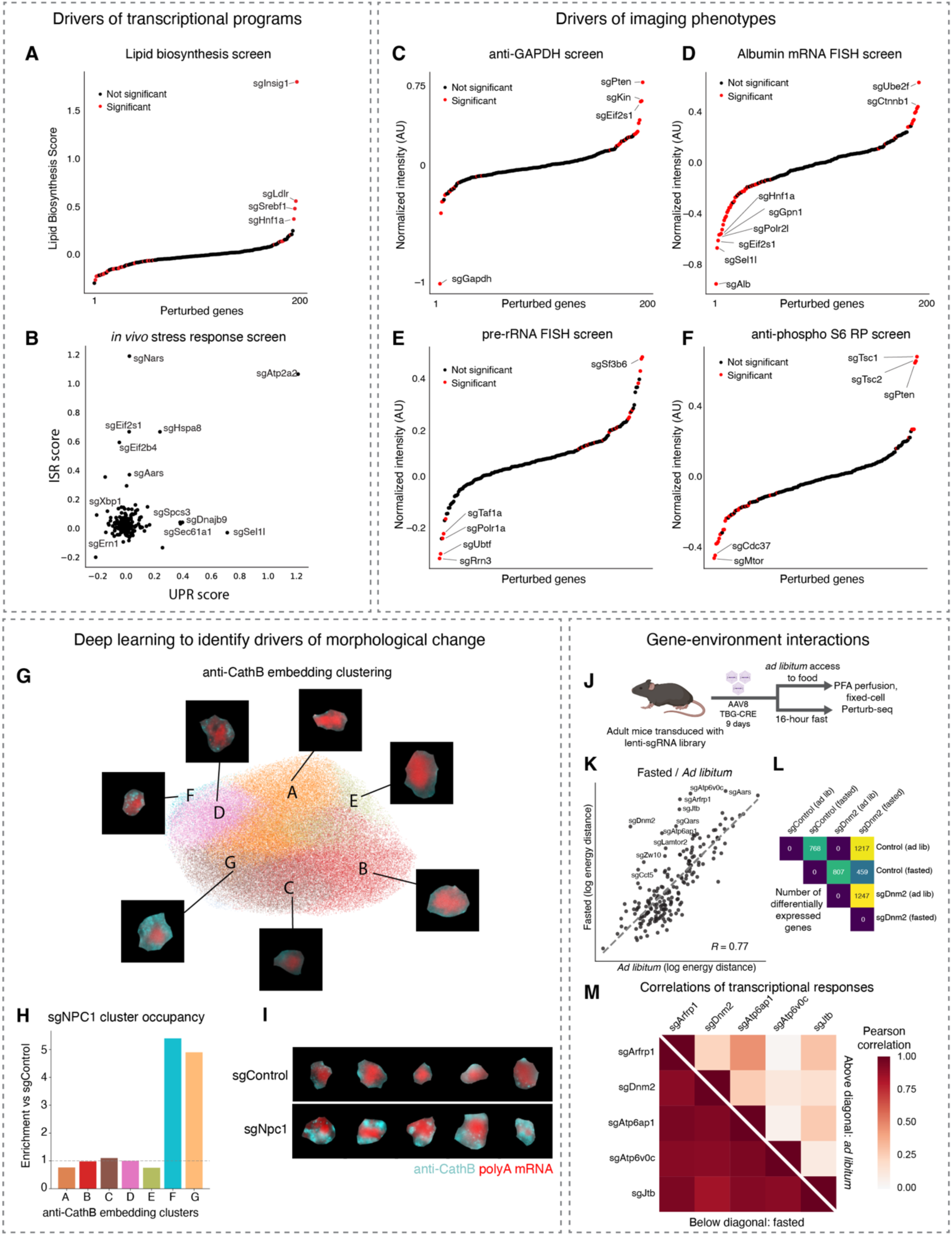
Identifying genetic drivers of liver physiology A. Ranking of perturbed genes by their impact on a literature-defined set of lipid and cholesterol biosynthesis genes in the Perturb-seq experiment. The genes included in the lipid score include *Hmgcs1*, *Sqle*, and *Fasn* and the score reflects mean log2-fold change across this panel, relative to negative control cells. Red reflects FDR-corrected significance (corrected p < 0.05, Benjamini-Yekutieli correction). B. Scatterplot showing the impact of perturbed genes on literature-defined sets of Unfolded Protein Response (UPR) and Integrated Stress Response (ISR) genes. The UPR score includes genes such as *Hspa5* and *Herpud1*; the ISR score includes genes such as *Atf4* and *Ddit4*. The scores are calculated relative to cells with control sgRNAs. C. Ranking of perturbed genes by their impact on anti-GAPDH intensity in the imaging experiment, relative to mean intensity in cells with control sgRNAs. Red reflects FDR-corrected significance (corrected p < 0.05, Benjamini-Yekutieli correction). D. Ranking of perturbed genes by their impact on *Albumin* mRNA FISH intensity in the imaging experiment, relative to mean intensity in cells with control sgRNAs. Red reflects FDR-corrected significance (corrected p < 0.05, Benjamini-Yekutieli correction). E. Ranking of perturbed genes by their impact on pre-rRNA FISH intensity in the imaging experiment, relative to mean intensity in cells with control sgRNAs. Red reflects FDR-corrected significance (corrected p < 0.05, Benjamini-Yekutieli correction). F. Ranking of perturbed genes by their impact on anti-Phospho S6 ribosomal protein intensity in the imaging experiment, relative to mean intensity in cells with control sgRNAs. Red reflects FDR-corrected significance (corrected p < 0.05, Benjamini-Yekutieli correction). G. A Leiden-clustered UMAP representation of CathB embeddings from the imaging experiment. Every point represents an individual cell. Sampled cells from each cluster are shown as insets. H. Bar plot of enrichment in each of the clusters in Npc1 knockout cells, relative to cells with control sgRNA. I. A sampling of cells with control sgRNAs or *Npc1* sgRNAs showing fluorescence micrographs the anti-CathB channel alongside polyA FISH. J. Schematic of the diet + genetic perturbation Perturb-seq experiment. Two diet conditions are tested, including food access (*ad lib*) and deprived (fasted) conditions. K. Scatterplot comparing the energy distance vs control cells for each knockout in the *ad lib* and fasted Perturb-seq datasets. The two distances are correlated (Pearson’s R = 0.77). L. Heatmap representing the number of differentially expressed genes between the indicated samples. Genes that are differentially expressed between the indicated conditions are defined with Benjamini-Hochberg-corrected, Mann-Whitney corrected p < 0.05. Cells are subsampled such that each of the comparisons includes the same number of cells. M. Heatmap representing Pearson correlations of pseudobulk transcriptional responses between the indicated knockouts, in the indicated condition. The knockout-specific transcriptional responses are calculated relative to cells with control sgRNAs from the same mouse. The phenotypes of these knockouts are more correlated in fasted tissue (mean Pearson’s R = 0.91, vs 0.19 in an *ad libitum* animal).

We also identified perturbations that activated transcriptional stress response pathways such as the integrated stress response (ISR) ^73^ or the unfolded protein response (UPR) ^74^, in hepatocytes in an intact liver (Figure 5B). We found that these two pathways were activated by the knockouts of separate sets of genes, suggesting that the stress-specific and selective transcriptional responses that are observed in cell culture are also a feature of hepatocyte physiology ^14,18,75^. For example, knockout of aa-tRNA synthetase genes such as *Nars* and *Aars* activated the ISR, as was previously observed in a genome-wide Perturb-seq experiment in cultured cells ^18^. Knockout of translation initiation factors such as Eif2α (*Eif2s1*) and Eif2b (*Eif2b4*) also activated the ISR. These results were consistent with established mechanisms of ISR activation by uncharged tRNAs and delayed translation initiation ^74^. Knockout of ER import and quality control genes such as *Sec61a1*, *Dnajb9*, and *Sel1l* activated the unfolded protein response. Knockout of the UPR effectors *Xbp1* and *Ire1* (*Ern1*) caused a decrease in UPR gene expression, suggesting some basal level of UPR activation in hepatocytes in *ad libitum* mice. Intriguingly, knockout of the ER calcium transport gene *Atp2a2* caused uniquely strong activation of both the UPR and the ISR, suggesting that calcium homeostasis plays a central role in regulating both ER function and broader cellular stress responses in hepatocytes.

For first-pass exploration of the imaging data, we ranked genetic perturbations by their impact on the intensity of each immunofluorescence or RNA FISH stain and identified perturbations that had a significant impact on the stain intensity. As a validation, we observed that sgRNAs targeting *Gapdh* had the strongest negative impact on anti-Gapdh staining intensity (Figure 5C) and sgRNAs targeting *Albumin* had the strongest negative impact on *Albumin* mRNA FISH staining intensity (Figure 5D). Targeting the RNA polymerase II machinery (*Polr2l*, *Gpn1*) or the core hepatocyte transcription factor *Hnf1a*, which directly binds the *Albumin* promoter, also had a strong negative impact on *Albumin* mRNA FISH, as did sgRNAs against the ER associated degradation (ERAD) factor *Sel1l*.

Interestingly, the screens correctly identified genetic regulators of many highly essential processes, suggesting that the ten days that we waited between AAV-Cre infection and tissue perfusion captured the acute impacts of perturbations. For example, targeting the RNA polymerase I machinery (*Polr1a*) and Pol I transcription factors (*Taf1a*, *Rrn3*, *Ubtf*) had the strongest negative impact on pre-rRNA FISH intensity (Figure 5E). Unexpectedly, targeting the nuclear speckle component *Sf3b6* caused an increase in pre-rRNA FISH intensity, suggesting a possible interplay between subnuclear structures (e.g. nuclear speckle and nucleolus) in hepatocytes *in vivo*.

Our screens also identified the *in vivo* regulators of cellular signaling pathways that play important roles in liver physiology. Phosphorylation of ribosomal protein S6 is a core function of the mTor signaling pathway. Knockout of *Mtor* and the potential mTor complex co-chaperone *Cdc37* caused the strongest decrease in anti-phospho S6 ribosomal protein intensity (Figure 5F) ^76,77^; knockout of the mTor inhibitors *Tsc1* and *Tsc2* and the growth inhibitor *Pten* caused the strongest increase in anti-phospho S6 ribosomal protein intensity ^77^. We note that imaging-based screening is uniquely powerful for probing genetic regulators of protein modification, which could not be measured by sequencing.

### Deep learning on the imaging-based perturbation data

In addition to modification of signaling proteins, imaging is also uniquely capable of detecting morphological changes beyond simple assessment of protein expression levels. We thus explored deep-learning-based approaches to explore more complex morphological phenotypes in the perturbation-imaging data. We used our autoencoder model to generate a lower-dimensional embedding for each image and then performed unbiased clustering on the embeddings. We illustrated these embeddings and the clustering with UMAP, using the imaging data of the lysosomal protein CathB as an example (Figure 5G). We then identified perturbations that shifted the representation of cells between the different clusters. For example, knockout of the lysosomal cholesterol transport gene *Npc1* caused significant enrichment of CathB morphologies in clusters F and G, which have many distinct anti-CathB punctae, and depletion from clusters A and E, which have very few anti-CathB punctae (Figures 5H and 5I). *Npc1* knockout also caused a significant increase in anti-CathB intensity (mean increase = 0.53 AUs, FDR-corrected p < 10^-34^). This is consistent with the accumulation of dysfunctional lysosomes that occurs when lysosomal cholesterol export is blocked, such as in Niemann–Pick disease type C with *NPC1* or *NPC2* mutations in humans ^78^.

### Investigating the impact of perturbations in altered physiological states

A key ability of performing screens *in vivo* rather than *in vitro* is to identify perturbations that are sensitive to changes in organismal physiology and homeostasis. We fasted a mouse from the lenti-sgRNA infected cohort for 16 hours prior to perfusion and tissue preparation and conducted a parallel Perturb-seq experiment on both the fasted animal and *ad lib* animal (Figure 5J). We used an energy distance metric to quantify the impact of each gene knockout on the distribution of global transcriptional states. We measured the impact of each knockout relative to cells with control sgRNAs in the same animal. The overall magnitude of transcriptional phenotypic changes caused by each gene knockout was similar between fasted and *ad libitum* mice for most genes (Figure 5K, Pearson’s R = 0.77). However, we noticed that some genes involved in autophagy and the lysosome (*Atp6v0c*, *Atp6ap1*, *Lamtor2*) and the broader endomembrane system (*Dnm2*, *Arfrp1*, *Jtb*, *Zw10*) had especially strong phenotypes in fasted animals in comparison to *ad libitum* animals. Consistent with the energy distance analysis, knockout of these genes caused many more genes to be differentially expressed in fasted mice than in *ad libitum* mice, with *Dnm2* knockout shown as an example (Figure 5L). This may be due to a higher dependence on lysosomal function during fasting ^79^; the failure of autophagy and endocytic function during fasting may cause severe cellular distress.

We examined the transcriptional phenotypes associated with knockout of these lysosome and endomembrane genes in *ad libitum* and fasted mice. The transcriptional responses associated with these knockouts were much more similar in a fasted animal (mean Pearson’s *R* = 0.91, versus 0.19 in an *ad libitum* animal; Figure 5M), suggesting that these knockouts caused a convergent phenotype in some physiological states. More broadly this illustrates how the modular structure of *in vivo* genotype-phenotype maps can be re-wired by environmental conditions ^80^.

### Three case studies exploring liver physiology with multimodal screening

#### Hepatocyte Zonation

Zonated gene expression in hepatocytes contributes to the spatial division of liver function. This zonation is believed to be maintained by gradients of morphogens, oxygen, nutrients, and hormones, but the requirements for these different processes to maintain zonal gene expression in a hepatocyte in an adult mouse are unclear. We explored our multimodal screening to shed light on mechanisms underlying hepatocyte zonal identity maintenance and dynamics.

Wnt signaling is an established driver of hepatocyte zonation. Wnt ligand secretion from central vein endothelial cells promotes pericentral gene expression in hepatocytes; low Wnt in the periportal region contributes to the periportal gene expression program ^61,81^. We scored all cells in the Perturb-seq dataset according to zonated gene expression and looked at the distribution of zonal expression in cells with knockouts of central Wnt effectors and inhibitors (Figure 6A; see Methods for the definition of zonal gene expression score). Knockout of the Wnt transducer *Ctnnb1* (β-catenin) caused a major decrease in the fraction of cells with pericentral gene expression, consistent with the role of Wnt signaling in pericentral expression ^82,83^. Conversely, knockout of the Wnt inhibitor *Apc* caused a decrease in the population of hepatocytes with periportal gene expression ^84^.

**Figure 6:**
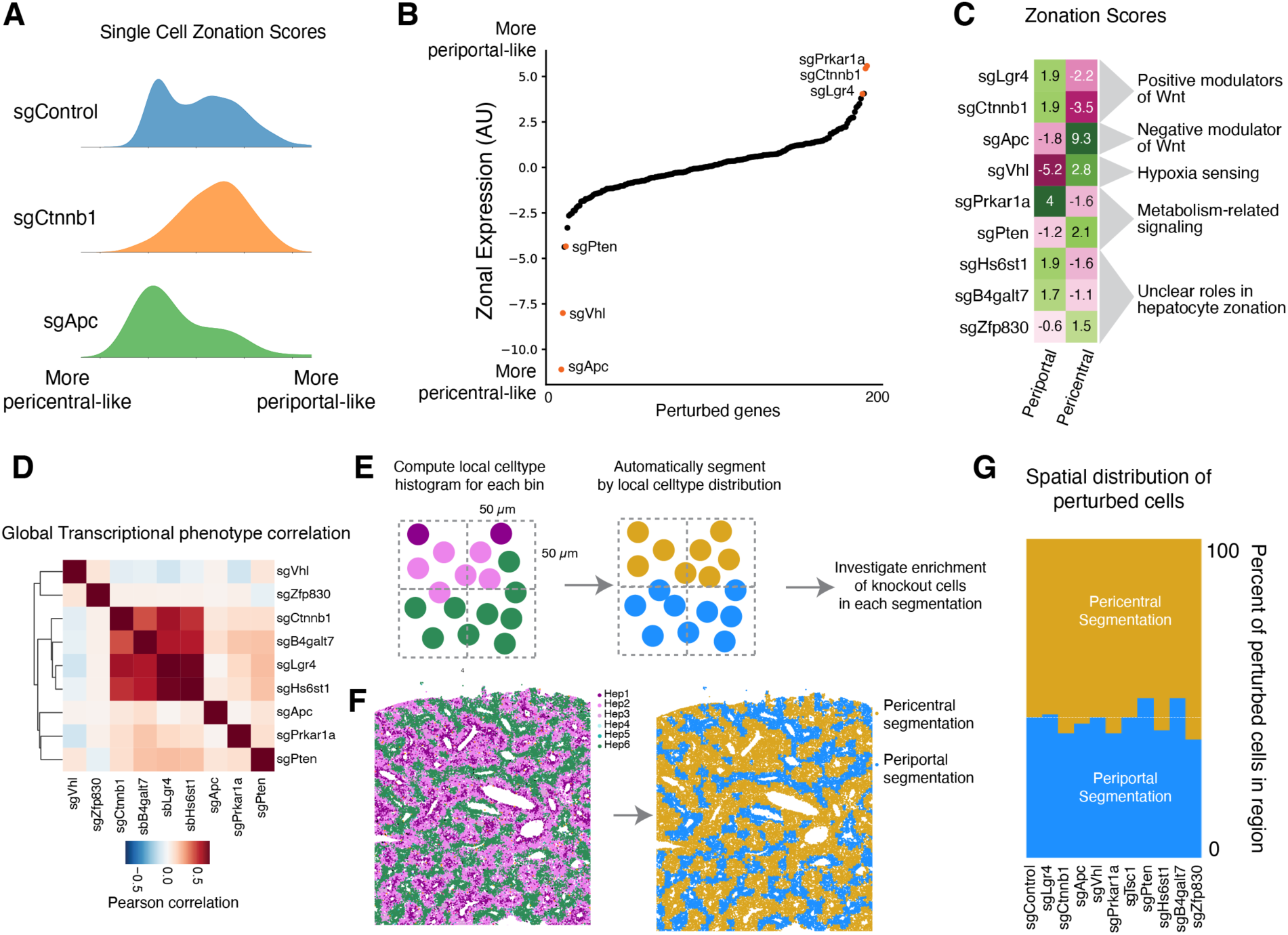
Multimodal investigation of the regulation of hepatocyte zonation A. Kernel density estimate plots showing the distribution of zonation gene expression in single cells with control sgRNAs, sgRNAs targeting *Ctnnb1*, and sgRNAs targeting *APC*. The single cell zonation scores reflect the expression of periportal marker genes such as *Cyp2f2* and *Hal* and pericentral marker genes such *Glul* and *Cyp2e1*, with periportal expression contributing positively and pericentral expression contributing negatively to the score. The scores are defined relative to the mean expression in cells with control sgRNAs. B. Ranking of perturbed genes by their average impact on the zonal gene expression score. C. Heatmap summarizing the categories of genes whose perturbation has a large impact on zonated gene expression. Here, the periportal and pericentral expression scores are shown separately. D. Heat map of the perturbation-perturbation correlation of pseudobulk transcriptional changes associated with the indicated sgRNAs. The color represents the Pearson correlation coefficient of pseudobulk phenotypes between indicated perturbations in the Perturb-seq dataset. E. Diagram of data-driven zonal segmentation process. The proportion of each of the different hepatocyte subtypes is calculated in 50 µm x 50 µm bins. The bins are then grouped into two zones based on the local cell type distribution and the enrichment of cells with each perturbation in the two zones is quantified. F. Representative tissue section showing cell types from the RCA-MERFISH experiment (left) and the resulting spatial segmentation of periportal and pericentral zones (right). G. Barplot of the fraction of cells in periportal and pericentral zones (as defined above), for the indicated genetic perturbations. The white line represents the fraction of cells with control sgRNAs. No perturbations have significantly altered spatial zonation.

We then looked across all gene knockouts in our experiment to identify the other drivers of zonal gene expression (Figure 6B). We also assessed periportal and pericentral expression as two separate scores, since the knockout of some core essential genes caused a general loss of zonal identity rather than a shift in one direction or the other (Figure 6C). Apc knockout caused the strongest increase in pericentral expression, as well as a substantial decrease in periportal expression. Knockout of *Vhl*, which mediates the degradation of hypoxia-inducible factors (HIFs) under normoxic conditions ^85^, caused a major decrease in periportal expression and also increased pericentral expression, consistent with the idea that oxygen tension directly regulates zonal gene expression ^61,86,87^. Loss of Vhl activity in the oxygen-rich periportal region may cause the specific decrease in periportal expression. Knockout of *Pten* and the gene *Zfp830*, which has not been studied in the context of hepatocyte zonation, also caused a shift toward a more pericentral-like expression state. Knockout of the R-spondin receptor *Lgr4* caused a strong increase in periportal gene expression, phenocopying *Ctnnb1*. Intriguingly, knockout of protein kinase a subunit *Prkar1a* also caused a substantial increase in periportal gene expression. Knockout of the heparan sulfate synthesis gene *Hs6st1* and the proteoglycan synthesis enzyme *B4galt7* also caused an increase in periportal expression and a decrease in pericentral expression, possibly by impacting the extracellular matrix in a manner that influences the interactions of signaling molecules such as Wnt ligands with their receptors ^88,89^.

Although one- and two-dimensional zonal expression scores described above are useful for exploration and interpretation of knockout effects on zonation, they do not necessarily reflect the full complexity of transcriptional phenotypes. Interestingly, when we compared the transcriptome-wide phenotype caused by these zonation-modulating knockouts, we found that knockout of some pro-pericentral factors, such as *Hs6st1* and *B4galt7*, closely phenocopied *Lgr4* and *Ctnnb1* knockout (Figure 6D), suggesting that their functions in hepatocytes are intertwined with Wnt signaling. However, knockout of the pro-pericentral factor *Prkar1a* caused a globally distinct transcriptional response despite its convergent impact on zonation-associated gene expression. Knockout of the apparent pro-periportal factors *Apc*, *Vhl*, *Pten*, and *Zfp830* all caused globally distinct transcriptional phenotypes despite their convergent impacts on zonal gene expression. These results emphasize the complexity of the relationship between cell state and metabolism in hepatocytes.

Our initial lentiviral transduction appears to target cells in all zones of the lobule; cells with control sgRNAs exhibit a full range of zonated gene expression (Figure 6A). In Perturb-seq, we measure the endpoint distribution of gene expression, but we do not know what happened during the time course of the perturbation or where each cell was in the tissue, prior to dissociation. What happens when a periportal hepatocyte gets a knockout such as sgApc that drives pericentral expression, or when a pericentral hepatocyte gets a pro-periportal sgRNA? The shift in endpoint zonal gene expression that we observe could be due to (1) *in situ* trans-differentiation without cell migration to an expression-appropriate zone, (2) trans-differentiation with migration, or (3) the death of cells in one zone and/or the proliferation of cells in the other zone.

To discriminate between these scenarios, we examined the perturbation-imaging data. We automatically segmented the tissue into two spatial zones (pericentral and periportal segmentations) and looked to see whether knockouts that impacted zonal gene expression also impacted the positions of cells with regard to the segmented pericentral and periportal zones (Figures 6E and 6F). Notably, the fraction of cells physically located in the pericentral and periportal zones did not change with any of the gene knockouts that significantly impacted zonal expression, even with potent knockouts such as *Apc* and *Ctnnb1* (Figure 6G). These results suggest that, on the time scale of our experiment and with our statistical power, cells with zonation-impacting knockouts exhibit gene expression changes due to trans-differentiation without migration or changes to survival or proliferation.

#### Stress Response

One major function of hepatocytes is to secrete abundant plasma proteins, so we were especially attuned to perturbations that impacted the endoplasmic reticulum. We noticed that knockout of the gene *Sel1l*, an ER-associated degradation (ERAD) co-factor that plays a crucial role in ER protein quality control ^90^, caused a strong increase in Calreticulin protein level (Figures 7A and 7C). This suggested that the transcriptional upregulation of UPR targets that we observed upon *Sel1l* knockout in the Perturb-seq experiment (Figure 5B) led to a corresponding increase in the ER protein chaperone capacity. *Sel1l* knockout also caused a significant decrease in *Albumin* mRNA level, but had relatively small impacts on the other proteins and abundant RNAs that we imaged (Figures 7B and 7C).

**Figure 7:**
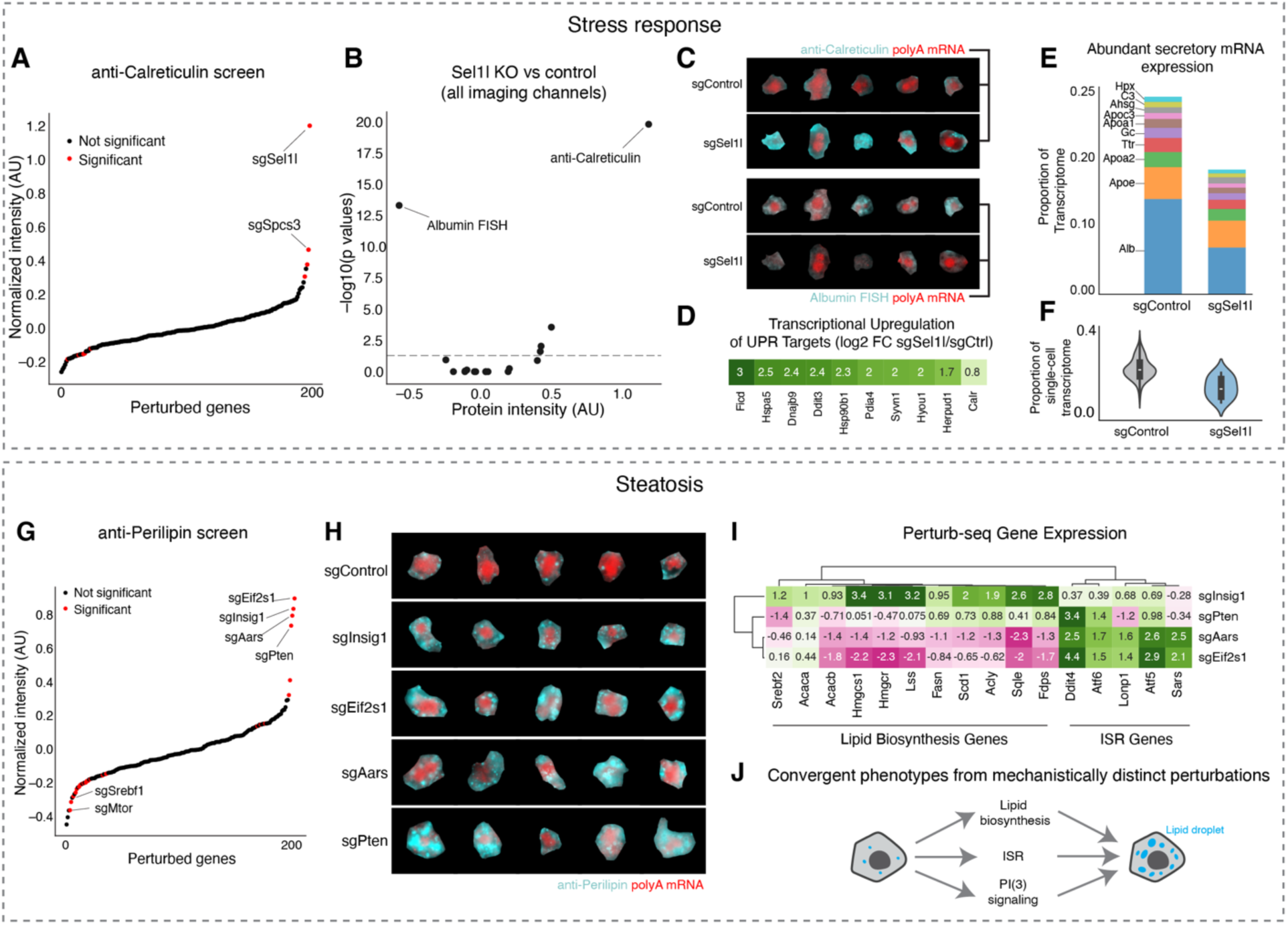
Multimodal investigation of stress response and steatosis A. Ranking of perturbed genes by their impact on anti-Calreticulin intensity in the imaging experiment, relative to mean intensity in cells with control sgRNAs. Red reflects FDR-corrected significance (corrected p < 0.05, Benjamini-Yekutieli correction). B. Volcano plot showing intensity change and significance of the imaged protein and RNA channels between cells with sgRNA targeting Sel1l vs cells with control sgRNAs. Dash line indicated corrected p value = 0.05. C. An unbiased sampling of cells with control sgRNAs and *Sel1l* sgRNAs, showing the anti-Calreticulin channel (top) and the *Albumin* mRNA FISH channel (bottom) alongside polyA FISH. D. Heatmaps showing log2-fold change in the expression of UPR genes in the sequencing experiment. E. Stacked bar plot representing the proportion of the measured pseudobulk transcriptome composed of the ten most abundant secretory mRNAs, ranked according to their abundance in control cells. F. Violin plot representing the fraction of each measured single-cell transcriptome that is composed of mRNAs encoding the ten abundant secretory mRNAs, from cells with sgRNAs targeting Sel1l and cells with control sgRNA. G. Ranking of perturbed genes by their impact on anti-Perilipin intensity in the imaging experiment, relative to mean intensity in cells with control sgRNAs. Red reflects FDR-corrected significance (corrected p < 0.05, Benjamini-Yekutieli correction). H. An unbiased sampling of cells with control sgRNAs and sgRNAs targeting the indicated gene, showing the anti-Perilipin channel alongside polyA FISH. I. A single-link hierarchically ordered heatmap showing log2 fold change in the expression of lipid biosynthesis genes and Integrated Stress Response genes, for the indicated genetic perturbations. J. Diagram illustrating convergent mechanisms that all cause lipid droplet accumulation. The knockouts may cause steatosis through three separate mechanisms: (1) through the activation of lipid biosynthesis in the case of Insig1 knockout; (2) through the sequestration of free lipids into lipid droplets alongside ISR activation, in the case of Eif2s1 and Aars knockout; and (3) a distinct, Pten-associated mechanism that may include update in plasma, lipid synthesis increase, and/or sequestration of free lipids.

Leveraging our paired imaging and sequencing data, we investigated the genome-wide transcriptional changes associated with *Sel1l* knockout in hepatocytes. As expected for an ER quality control factor, *Sel1l* knockout caused the upregulation of many other ER chaperone mRNAs, such as *Hspa5* and *Dnajb9*, as well as *Calreticulin* itself (Figure 7D). *Sel1l* knockout decreased *Albumin* mRNA by 50% (Figure 7E), as expected from our imaging data, but intriguingly it also caused the downregulation of other mRNAs that encode for a range of abundant secreted plasma proteins, such as Apolipoprotein AI (*Apoa1;* 38% decrease) and the GC Vitamin D Binding Protein (*Gc*, 36% decrease) (Figure 7E). Altogether, the mRNAs downregulated by *Sel1l* knockout reflected an ∼40% decrease in the total quantity of abundant secreted protein mRNAs (Figure 7F). These results suggested that a major function of the unfolded protein response in hepatocytes *in vivo* is to decrease the burden of secreted protein mRNAs and therefore of nascent proteins on the ER translocon and folding machineries. This decrease in mRNA abundance could occur indirectly through transcriptional downregulation ^91^ or directly through post-transcriptional stress response pathways such as Regulated Ire1a-Dependent Decay (RIDD), where it was found that ER stress sensor Ire1 can directly cleave mRNAs of secreted proteins on the ER surface to lower the flux of proteins through the ER^92,93^.

#### Lipid Accumulation

Hepatic steatosis, characterized by the accumulation of lipids within hepatocytes, is linked to metabolic syndrome and can progress to serious diseases such as metabolic dysfunction-associated steatohepatitis (MASH) and fibrosis ^94^. We used our imaging screen to identify *in vivo* drivers of steatosis. Acute knockout of the lipid biosynthesis inhibitor Insig1, the growth inhibitor Pten ^95^, the integrated stress response inhibitor Eif2α (*Eif2s1*), and the alanine-tRNA synthetase *Aars* all caused steatosis, as measured by enlargement and increase in signal from lipid droplets, marked by anti-Perilipin (Figures 7G and 7H).

We then examined the Perturb-seq data from the same mice to identify the transcriptome-wide changes associated with each of these four perturbations (Figure 7I). Consistent with its function as an inhibitor of lipid biosynthesis gene transcription ^96^, *Insig1* knockout caused the upregulation of mRNAs for cholesterol biosynthesis genes such as *Hmgcr* and *Fdps* and fatty acid biosynthesis genes such as *Acaca* and *Fasn*. On the other hand, knockout of *Aars* and *Eif2s1* drove upregulation of ISR target genes such as *Ddit4* and *Atf5*. Intriguingly, *Aars* and *Eif2s1* knockout actually caused a decrease in most lipid biosynthesis mRNAs. Finally, knockout of *Pten* caused yet another distinct transcriptional response that could not be summarized either as the ISR or lipid biosynthesis.

Thus, through integrated analysis of Perturb-seq and imaging, we found that knockouts of these four genes apparently drove three separate transcriptional responses but converged on the steatotic phenotype. Considering these results together, we propose that the knockouts cause steatosis through three separate mechanisms—(1) through the transcriptional activation of lipid biosynthesis genes in the case of *Insig1* knockout; (2) through the apparent sequestration of free lipids into lipid droplets in the case of *Eif2s1*/*Aars* knockout, possibly as a component of the ISR that may serve to protect the cytoplasm and other organelles in stressed cells from biophysically disruptive lipid molecules ^97–99^, and (3) a distinct *Pten*-associated mechanism, possibly including uptake from plasma, some synthesis increase, sequestration, and/or the repurposing of organellar stores ^100–103^ (Figure 7J).

## DISCUSSION

Massively parallel *in vivo* genetic screens provide a powerful approach to dissect regulators of cellular and tissue physiology in their native context ^38^. Here, through extensive technical development and integration of multiple single cell profiling methods, we establish Perturb-Multi, a general and scalable approach for multimodal, pooled genetic screens with Perturb-seq- and multiplexed-imaging-based phenotyping. This multimodal *in vivo* approach allows us to access and interrogate physiological processes that are difficult to study by other means. We conduct screens in hepatocytes in the mouse liver and demonstrate how the resulting data enables causal dissection of multifarious aspects of liver biology, from hepatocyte zonation to steatosis to the signaling of nutritional status. Our data also provide a rich resource for future studies of liver biology and development of computational methods. In particular, our multimodal imaging data should prove useful for developing new analytical and machine learning tools for dimensionality reduction, feature extraction, and cross-modal analysis. More generally, this approach will enable the creation of large-scale training datasets for machine learning and AI efforts to build models of “virtual” cells, driving both new discoveries and new forms of understanding ^37^.

We show that the combination of transcriptomic profiling and subcellular morphology imaging enables new analyses of cellular heterogeneity within tissue, under both homeostatic and perturbed physiological conditions. Tissue-scale spatial analysis of protein feature embeddings revealed zonal heterogeneity in several imaging targets, beyond the expected *Albumin* mRNA and Perilipin. Cross modal analysis of subcellular protein and abundant RNA morphology across transcriptionally-defined hepatocyte subtypes revealed a continuum of morphological types between different liver zones. More generally, the multimodal combination of spatial transcriptomics to define cell types and states with morphological features that can be linked to the extensive histopathology literature should enable studies of both healthy and diseased tissue.

Our screening results demonstrate the value of multimodal phenotyping that combines both multiplexed protein and RNA imaging and transcriptome-wide mRNA sequencing to study genetic regulators of cellular state *in vivo*. Applying this approach to cells in native tissue allows us to obtain insights into three core aspects of liver physiology in a single experiment: hepatocyte zonation, dynamic stress responses, and lipid droplet accumulation.

Zonation: Hepatocytes in different zones of the liver express different genes and have different functions^61^. The regulation of hepatocyte zonation has been challenging to study *in vitro*, as hepatocytes rapidly lose their zonal identity when isolated and introduced into cell culture. Even complex organ-on-a-chip and organoid models provide imperfect recapitulations of tissue architecture ^104,105^. Here, we directly probe the regulation of zonal gene expression in the mouse liver. In addition to confirming the known roles of oxygen sensing and Wnt signaling in regulating zonal identity ^61^, we show that even brief (<10 day) disruption of these pathways rewires hepatocyte zonal gene expression in adult mice, apparently in a cell autonomous manner. Remarkably, we further identify several novel candidate regulators of zonation such as *Hs6st1* and *B4galt7* that have an equal impact on zonation relative to perturbing the Wnt pathway, illustrating the value of systematic exploration. Notably, these factors are involved in the modification of secreted and extracellular matrix proteins and may point toward a unique capacity of *in vivo* screens to explore causal relationships between the extracellular microenvironment and intercellular signaling. By analyzing the spatial location of the genetically perturbed cells, we show that upon deletion of these key regulatory genes, cells change their transcriptional zonal states without physical relocation across pericentral and periportal zones in the tissue. We anticipate future experiments with genome-scale perturbation libraries and multiplex perturbation vectors to comprehensively classify zonation regulators, and to identify epistatic relationships between the associated signaling pathways.

Stress response: Hepatocytes are secretory cells that must produce Albumin, among other proteins, in massive quantities. As such, stress response pathways that monitor protein folding in the endoplasmic reticulum, such as unfolded protein response (UPR), play an important role in maintaining hepatocyte function. Notably the UPR is activated in several liver diseases, including viral hepatitis, alcohol-associated liver disease, and MASH ^106,107^. The secretion of proteins such as Albumin drops dramatically when the liver is dissociated and hepatocytes are introduced into an *in vitro* setting ^108^, indicating the importance of *in vivo* models for the investigation of hepatocyte secretion. In our imaging-based pooled genetic screening, we found that the knockout of *Sel1l*, an important ubiquitin ligase adaptor protein involved in ER-associated degradation, caused massive upregulation of the ER quality control protein Calreticulin, and downregulation of *Albumin* RNA. In subsequent analysis of the Perturb-seq data, we found that Sel1l knockout caused both transcriptional upregulation of UPR target genes and the 40% transcriptional downregulation of abundant secreted proteins. We thus hypothesize that a major function of the UPR in hepatocytes in the liver is to downregulate secretory protein genes, thereby reducing the burden on the ER folding machinery, either transcriptionally or through post-transcriptional mechanisms such as RIDD ^93^. The extent to which UPR-associated down-regulation of secreted proteins may contribute to hypoproteinemia seen in chronic liver disease remains an outstanding question.

Lipid droplets: Lipid droplets play a crucial role in hepatocyte energy storage and lipid metabolism. Accumulation of lipid droplets is key to the pathology of metabolic dysfunction-associated steatotic liver disease (MASLD), whereby progress from steatosis to steatohepatitis is marked by distinct transcriptional states in humans ^109^. As an example of interrogating *in vivo* physiology, we find that genetic knockouts targeting *Insig1*, *Eif2s1*/*Aars* and *Pten* all caused a convergent morphological phenotype with the dramatic accumulation of lipid droplets, but entirely distinct transcriptional responses. While the highly interpretable imaging results indicate that the function of these genes impinges on lipid homeostasis, their distinct effects on the transcriptional states indicate they likely induce lipid accumulation through distinct mechanisms. Our paired imaging and sequencing approach was necessary to reach this conclusion. Imaging lipid droplet accumulation on its own would reveal the ultimate cellular consequences of the perturbations; measuring the transcriptional responses by Perturb-seq would show the complexity and distinctiveness of each perturbation, but mask their common functional impact on the cell.

These insights into hepatocyte biology suggest that applying our approach to other organs will be of immediate value. The basic approach that we have developed here is applicable to dissociated cells or sections from any tissue. Similar genetic mosaics can be created in other organs using viral delivery, including the brain and skin ^29–31^, although there are currently technical challenges to achieving the same level of uniformity and precision of delivery as in the liver. In particular, AAV-based delivery will further expand the number of accessible organs to include heart, lung, brain, and skeletal muscle – potentially even spanning the majority of tissues in an animal. AAV-based approaches can also be applied in non-murine species, including non-human primates. AAV delivery of sgRNAs has recently been used for Perturb-seq in the mouse brain ^29,30^; we anticipate that it can be adapted to our platform of paired imaging and fixed-cell Perturb-seq. AAV delivery can suffer from problems of cell type tropism and uncontrolled viral multiplicity-of-infection (MOI), which may make pooled screens more challenging to interpret. Future multimodal mosaic screens will build on advances in technology for the delivery of perturbation reagents, such as engineered AAVs that target specific cell types with high efficiency ^110^, or methods to create controllable MOI. Tissues more complex than the liver will provide new challenges and opportunities for understanding gene function across a variety of cell types and states.

Although our experiments demonstrate the power of multimodal functional interrogation of diverse genes *in vivo* at the scale of hundreds of genes, our approach is highly scalable in terms of the number of phenotypes, genotypes, and perturbed cells measured. In terms of phenotype, the number of genes imaged using MERFISH ^36,111,112^, as well as the number of proteins assayed through multiplexed immunofluorescence, could be readily increased in future experiments to obtain a more unbiased picture of the phenotypic states of cells. In terms of genotype, a clear future direction is to increase the number of perturbations, ultimately to the genome-wide scale, enabling truly unbiased and comprehensive mapping of gene function.

Achieving genome-wide scale will require measurements of more individual cells. The number of cells is not limiting: each mouse liver contains 10^8^ hepatocytes, enabling millions of perturbed cells per experiment. While the time and/or cost spent on an experiment scales linearly with the number of perturbations measured, the value of the information collected could scale superlinearly - this is especially true for investigating the interactions and relationships between different genotypes and phenotypes, allowing different genes to be grouped into pathways or complexes ^18,24^. Increasing the number of cells measured per perturbation will also allow us to resolve more subtle phenotypes, as well tackle more complex and heterogeneous tissues containing larger numbers of distinct cell types or cell states. Achieving larger scales through sequencing is currently primarily limited by cost, which could be reduced by superloading microfluidic droplets ^113^ or using scalable split-pool approaches ^114,115^. While imaging is already quite high-throughput – scales of tens to hundreds of thousands of cells per day ^116^ and millions of cells per experiment are possible ^117^ – reaching genome-wide scale will require further improvements in microscopy instrumentation and analysis to increase the throughput (for example, ∼20,000 perturbations x ∼100 cells per perturbation = ∼2 million perturbed cells, which requires ∼20 million imaged cells when probed at a low MOI (∼0.1) to ensure most perturbed cells harboring a single perturbation). For both imaging and sequencing, throughput could be further increased by infecting cells at higher MOI then deconvolving phenotypes, using ideas from compressed sensing ^118^.

These large-scale, multimodal maps will enable both biological discovery as well as the training of sophisticated AI models that can capture the rich structure of genetic and cell function. In particular, these data will provide fuel for emerging large-scale “virtual cell” efforts that build on existing cell atlases to model cellular state across diverse cell types, tissues, and organisms ^119–121^. Such generative models promise to enable vastly better prediction of the effects of perturbations on cells, with applications ranging from reverting disease phenotypes to enabling entirely new functionalities ^38^. Genome-wide Perturb-seq data has already been successfully applied to build predictive models of the effects of perturbations on cells ^122^. Such models are currently constrained by the paucity of available perturbation data and the limited phenotype modalities measured, requiring more perturbations and richer phenotypes in diverse cell types to generalize beyond cultured cells. Adding an understanding of causality to these models through training on large-scale perturbation data will dramatically increase their predictive power, enabling new approaches to discovery and hypothesis generation, cellular engineering, and therapeutic development.

### Limitations of the study

Some aspects of our study limited the scope of our results. (1) We targeted only ∼200 genes that are potentially important for liver physiology, limiting the number of pairwise comparisons that could be made between knockouts and the chance for serendipitous discoveries of novel gene functions, (2) we delivered perturbations only to hepatocytes and sampled a limited number of cells per perturbation, limiting our ability to understand how the phenotypic impacts of knockouts vary across cell types in the liver, as well as to study non-cell-autonomous effects, with high statistical power, and (3) our immunofluorescence panel covers only a subset of proteins, cellular structures, and cell signaling pathways, limiting the number of morphological phenotypes measured.

## Data and Code Availability

Sequencing data have been deposited with GEO (GSE275483). Cell images will be deposited to FigShare. Our codebase for the RCA-MERFISH pipeline and perturbation analysis will be available on GitHub: https://github.com/weallen/InVivoMultimodalPerturbation

## Acknowledgements

We thank K. Loh, L. Luo, M. Lovett-Barron, P. Rajasethupathy, S. Konermann, P. Hsu, D. Lara-Astiaso, F. Zhang, X. Wang, E. Richman, K. Knouse, H. Keys, K. Smolyar, L. Koblan, G. Muthukumar, B. Herken, M. Schnall-Levin, P. Smibert, J. Durruthy, A. Kohlway, P. Lund, R. Sauer, T. Norman, D. Ron, C. Gross, and all members of the Allen, Weissman, and Zhuang labs for helpful discussions. We thank C. Muresan and N. White for administrative support; S. Sinha for laboratory support; the Whitehead GTC and S. Gupta for support with sequencing; the Whitehead Flow Cytometry Core and P. Autissier for support with FACS; and A. Halpern for support with imaging.

This research was supported by:

- Howard Hughes Medical Institute (C.D., J.S.W., X.Z.)
- Star-Friedman Challenge Award (W.E.A., X.Z.)
- Milton Fund Award (W.E.A.)
- Longevity Impetus Grants/Hevolution Foundation (W.E.A., X.Z.)
- National Institutes of Health (NIH) Centers of Excellence in Genomic Science (2RM1HG009490-07) (J.S.W)
- Chan Zuckerberg Initiative (J.S.W.)
- Arc Institute (W.E.A.)
- Gates Foundation (X.Z.)
- Whitehead Innovation Initiative (J.S.W.)
- The Ludwig Center at MIT (J.S.W.)
- Fannie and John Hertz Foundation Fellowship (R.A.S.)
- NSF Graduate Research Fellowship (R.A.S.)
- Harvard Society of Fellows (W.E.A.)
- National Institute of Health T32 3T32DK007191-50S1 (J.S.)
- Burroughs Wellcome Fund Career Award at the Scientific Interface (W.E.A.)
- Jane Coffin Childs - HHMI Fellowship (X.P.)
- National Institute of Health T32 3T32DK007191-50S1 (J.S.)

## Author Contributions

R.A.S. and W.E.A. jointly developed the project’s methodology and experimental design with J.S.W. and X.Z.. R.A.S. and W.E.A. performed all experiments, with help from J.L.. R.A.S., W.E.A., and X.P. performed analysis and visualization, with help from T.K.L.. X.P. led deep learning efforts. J.S. contributed expertise in liver biology and conducted preliminary fixed-tissue dissociation experiments with R.A.S.. Z.A.S., R.A.S., and W.E.A. performed high-fat diet and fasting experiments. C.D. contributed animal resources and guidance. R.A.S., W.E.A., X.P., J.S.W., and X.Z. interpreted data. R.A.S, W.E.A., J.S.W. and X.Z., wrote the original manuscript, and all authors reviewed and edited it. W.E.A. initiated this collaboration between the X.Z. and J.S.W. labs, as an independent Junior Fellow working in the laboratories of X.Z. and C.D. at Harvard. W.E.A., J.S.W., and X.Z. led and supervised the project.

## Methods

### RESOURCE AVAILABILITY

Lead Contact: Further information and requests for resources and reagents should be directed to and will be fulfilled by the lead contact.

### EXPERIMENTAL MODEL AND STUDY PARTICIPANT DETAILS

#### Animals

Male and female wildtype and B6;129-Gt(ROSA)26Sortm1(CAG-cas9*,-EGFP)Fezh/J in a C57BL/6J background were used in this study. Mice were obtained from Jackson labs or bred at Harvard or MIT. Mice were maintained on a 12 hr light/ 12 hr dark cycle (14:200 to 02:00 dark period) at a temperature of 22 ± 1oC, a humidity of 30-70%, with *ad libitum* access to food and water unless otherwise noted. Mice were fed *ad libitum* with a diet containing 60% kcal from fat (Research Diets, Inc. D12492i) or standard chow. Animal care and experiments were carried out in accordance with NIH guidelines and were approved by the Harvard University and Whitehead Institute Institutional Animal Care and Use Committees (IACUC).

#### Cell lines

HEK 293T/17 (ATCC CRL-11268) cells were cultured in DMEM supplemented with glutamax, HEPES, 10% fetal bovine serum, 100 units/mL penicillin, and 100 mg/mL streptomycin (ThermoFisher Scientific). K562-CRISPRi cells (Gilbert et al, 2014) were cultured in RPMI supplemented with glutamine, HEPES, 10% fetal bovine serum, 100 units/mL penicillin, and 100 mg/mL streptomycin (ThermoFisher Scientific). AML12 cells (ATCC CRL-22540 were cultured in DMEM/F12 medium supplemented with 10% fetal bovine serum, 10 mg/mL insulin, 5.5 mg/mL transferrin, 5 ng/mL selenium, and 40 ng/mL dexamethasone (ThermoFisher Scientific).

### METHOD DETAILS

#### Lentiviral Construct Cloning

The parental mosaic screening vector (pVV1) was generated from a modified CROP-seq vector (pBA950; addgene 122239 ^123^) and from a lentiviral vector used for CRISPR screens in the liver (pLentiCRISPRv2-Stuffer-HepmTurquoise2; addgene 192826 ^27^) using standard methods. First, the EF1a-BFP was replaced by mTurquoise driven by a hepatocyte-specific promoter. Then, the parental mU6 promoter was replaced by an mU6 promoter that did not contain a loxP site ^124^, yielding pVV1.

#### CRISPR Guide Library Cloning

The parental pVV1 vector was digested with BstXI and BamHI and purified by gel electrophoresis. Library elements containing both sgRNAs and their associated barcodes were ordered as eblocks and pooled before cloning. The library elements were synthesized with three different sgRNA constant regions, which decreases recombination between the sgRNA and the barcode during lentiviral packaging^14^. The library elements were also digested with BstXI and BamHI and purified by gel electrophoresis. The digested library was ligated into digested pVV1. The ligation was purified with a silica column (Zymo) and electroporated into MegaX DH10B T1 R electrocompetent cells (ThermoFisher Scientific) according to the manufacturer’s protocol. The cells were recovered and then directly introduced into liquid culture and maxiprepped the next day. Samples were plated and used to confirm >1000x library coverage during cloning. Colonies were sequenced to confirm cloning fidelity.

#### Lentiviral Preparation

The lentivirus was generated according to standard methods by the transfection of 293T/17 cells with Fugene HD (Promega), our library vector, psPAX2, and pMD2.G. We used ViralBoost (Alstem) according to the manufacturer’s instructions. We harvested supernatant, filtered it through a 0.45 µm PES membrane, and conducted a 10x concentration with PEG and NaCl (Lenti-X Concentrator; Takara) according to the manufacturer’s instructions. We then pelleted this concentrate by centrifugation in a swinging bucket centrifuge (25,000*g*, 2 hours, 4°, Beckman Coulter SW 32 Ti) and resuspended it in cold PBS + 4% glucose, leading to an additional 100x concentration. Lentivirus was flash frozen and titered approximately through the transduction of AML12 cells.

#### Mosaic Liver Preparation

The lentivirus was thawed on ice and up to 50ul was injected into the temporal vein of postnatal day one mice ^27,125^. We injected approximately 1 x 10^7^ titer units of lentivirus per animal. We then allowed the mice to grow to adulthood (>P30) and induced Cas9 through the retro-orbital injection of AAV8 with Cre driven by a hepatocyte promoter (Addgene 107787-AAV8; ∼5 x 10^11^ genome copies per animal). We maintained the mice for ten days, or alternatively maintained for nine days and then fasted for 16 hours. The mice were anesthetized and fixed by perfusion of PBS + 4% paraformaldehyde. Livers were removed and fixation was continued for 3 hours in PBS + 4% paraformaldehyde, followed by ∼13 hours in PBS + 4% paraformaldehyde + 30% sucrose. Fixed livers were then frozen in Optimal Cutting Temperature compound (OCT) and stored at -80°.

#### Mosaic Liver Dissociation for Fixed Cell scRNA-seq

100µm sections of liver were generated on a cryostat and stored at -80° for later processing. The sections were washed with cold 0.5x PBS to remove residual OCT and then resuspended in warm RPMI + 1 mg/ml Liberase Th. The material was transferred to a gentleMACS C tube and dissociated on a gentleMACS Octo Dissociator with heaters (Miltenyi) with the program 37C_FFPE_1. The cells were strained through a 30µm pre-separation filters (Miltenyi) and singlets (diet experiment) or GFP+, mTurqiouse+ singlets (Perturb-seq experiment) were isolated by FACS (ARIA II, BD) and maintained at 4° in 0.5x PBS until scRNA-seq.

#### sgRNA probes for Fixed Cell scRNA-seq

Probes for the sgRNAs were obtained from IDT as opools. Probe sequences are found in Table S9. The left-hand side probes targeted the sgRNA constant region; three variants of this probe were included in each hybridization, targeting each of the three constant regions included in the sgRNA library. The left-hand side probes also had a sample barcode sequence. The right hand side probes were 5’ phosphorylated and targeted the protospacer sequence directly. The barcode spike in probes contained the appropriate TruSeq sequences such that they could be amplified separately from the transcriptome probes and sequenced independently. We included a variable number of N bases to ensure base diversity during sequencing.

#### Fixed Cell scRNA-seq Library Preparation and Sequencing

scRNA-seq was conducted with the Single Cell Gene Expression Flex platform (10x Genomics). Spike-in probes for the sgRNAs were included to a final concentration of 2 nM. Cells were counted on a Countess II (ThermoFisher Scientific). We used four sample barcodes and recovered the cells across one (diet experiment) or eight (Perturb-seq experiment) of microfluidic channels. mRNA libraries were prepared according to the manufacturer’s instructions. sgRNA libraries were prepared with the Fixed RNA Feature Barcode Kit according to the manufacturer’s instructions. Libraries were sequenced using a NovaSeq 6000 (Illumina).

#### scRNA-seq Alignment and Calling

The scRNA-seq mRNA data was aligned with CellRanger (10x Genomics). The sgRNA reads were aligned with a custom pipeline using the cell barcodes produced by CellRanger. Briefly, we used bowtie2 (flags --very-sensitive --local) to align the reads to the sgRNA probe library. We then selected sgRNA reads with a cell barcode that was shared with a cellranger-called cell and identified the number of UMIs for each sgRNA in each cell. We tested several perturbation calling approaches including mixed model calling and identifying outliers in a Poisson distribution or zero-inflated Poisson distribution and found the best performance across sgRNAs with thresholding, choosing thresholds empirically to maximize on target knockdown of perturbation targets with apparent NMD. We identified all cells with exactly one sgRNA over the chosen threshold and excluded the rest from all analyses.

#### Antibody Labeling

Antibodies were obtained in >50 ug quantities and labeled with bifunctional 5’ Acrydite - bit sequence - 3’ DBCO oligos (IDT) by enzymatic modification and click chemistry (SiteClick Antibody Azido Modification Kit, ThermoFisher) according to the manufacturer’s instructions. Antibody-oligo conjugates were concentrated in PBS by ultrafiltration with a 100kDa membrane (Millipore), which also removed residual un-conjugated oligonucleotides. Antibody-oligo conjugates were aliquoted into tube strips and snap frozen in liquid nitrogen.

#### RCA-MERFISH Readout Probe Synthesis

Amine-modified 15mer oligonucleotides were obtained from IDT (standard desalting). Oligos were resuspended to 300 µm in 112 mM sodium bicarbonate solution (ThermoFisher). 300 mM Sulfo-Cy3-NHS ester, Sulfo-Cy5-NHS ester, and Sulfo-Cy7-NHS ester (Lumiprobe) solutions were made in dry DMSO (Sigma Aldrich). The appropriate dye was added to each oligo to a final concentration of 10 mM and the dyes were allowed to react for 24 hours in the dark at room temperature. Sodium acetate pH 5.5 was added to a final concentration of 500 mM and then ice cold ethanol was added to a final concentration of 80%. The oligos were incubated at -20° C for >24 hours, pelleted by centrifugation at >18,000*g* at 4° C for >20 minutes, washed 3x with ice cold 80% ethanol, dried in a vacufuge, and resuspended in TE pH 8 to a final concentration of 100 µm. Labeled oligos were stored at 4° C until use.

#### RCA-MERFISH Encoding Probe Design and Construct

For the 205 endogenous genes, we used a 21-bit, Hamming Weight 4, Hamming Distance 4 codebook for MERFISH imaging. For the 456 perturbation barcodes, we used an 18-bit, Hamming Weight 6, Hamming Distance 4 codebook for MERFISH imaging. For both the endogenous genes and perturbation barcodes, individual genes/barcodes were randomly assigned to codewords in the codebook. For each gene/barcode, we designed a total of 8 (endogenous genes) or 3 (barcodes) encoding probes targeting the gene (Table S2) or barcode (Table S7) mRNA sequence. Following the guidelines for MERFISH probe design as previously described ^126^, we selected 60 mer regions that could be split into two 30 mers, where each half had GC content between 30 and 70%, melting temperature Tm within 60-80oC, isoform specificity index between 0.7 and 1, gene specificity index between 0.75 and 1, and no homology longer than 15 nt to rRNAs or tRNAs. Pairs of adjacent probes that had a ligation junction with a G or C at the donor (5’ phosphorylated) end of the probe were excluded. The two halves of the probes were then split, and between them was added an RCA primer sequence and the reverse complements of the 4 (endogenous) or 6 (barcode) readout sequences that encoded the identity of that gene. PCR handles with BciVI (left hand side) and BccI (right hand side) restriction sites were then appended to either end of the probe. The readout sequences on the encoding probes are detected with dye-labeled readout probes with complementary sequences in order to decode the gene or barcode.

#### RCA-MERFISH Encoding Probe Synthesis

RCA-MERFISH encoding probe libraries were first synthesized at femtomolar scale in a pool by Twist Biosciences. Probe libraries were then amplified by limited-cycle PCR (Phusion Polymerase, New England Biolabs), purified using SPRI beads (Beckman Coulter), and then blunt-end ligated using T4 DNA to circularize. Circularized DNA molecules were nicked using Nt.BbvCI (NEB), and then RCA amplified overnight at 30oC using Phi29 DNA polymerase from the nick site with T4 gene 32 single-stranded binding protein (SSB) (New England Biolabs). RCA amplified DNA was ethanol precipitated and resuspended in CutSmart buffer. Oligos with degenerate ends containing restriction enzyme sites were annealed to the RCA product, and the mixture was digested overnight with BccI and BciVI (New England Biolabs). The final libraries were then purified using magnetic beads, eluting in Tris-EDTA (TE) (ThermoFisher) buffer to a final concentration of ∼10 nM/probe. The resulting library was stored at – 20oC until use.

#### RCA-MERFISH Sample Preparation

Sample preparation occurred over several days. Blocking, antibody staining, MelphaX modification, probe hybridization, ligation, and RCA were conducted with the coverslips inverted onto small volumes of reaction mixture over parafilm, whereas decrosslinking, washing, and digestion were conducted with the coverslips upright in >5 ml of solution in 60 mm tissue culture dishes. Silanized, PDL-coated coverslips were prepared according to the method of ^43^.

10um sections of liver were cut onto silanized, PDL-coated coverslips, warmed to room temperature for 15 minutes, and attached to the surface by 15 minutes of postfixation in PBS + 4% paraformaldehyde. The sections were washed 3x with PBS and decrosslinked at 60° in TE pH 9 (Genemed) for an hour. The sections were then washed with PBS.

The sections were blocked at RT for 20 minutes in blocking buffer (1x PBS, 10 mg/ml BSA (UltraPure, ThermoFisher), 0.3% Triton X-100, 0.5 mg/ml sheared salmon sperm DNA (ThermoFisher), and 0.1 U/µL SUPERase·In RNase Inhibitor (ThermoFisher) ^127^. It was crucial that the blocking buffer did *not* contain dextran sulfate, as even trace amounts inhibited later ligation and/or RCA.

The antibody-oligo conjugates were pooled and diluted into blocking buffer. The sections were then stained overnight at 4° with this mix. We estimate that we stained each coverslip with ∼150 ng of each oligo-antibody, diluted into 150 ul of blocking buffer. The sections were then washed 3x with PBS and then incubated with PBS at RT for 15 minutes, then postfixed for 5 minutes in PBS + 4% paraformaldehyde. The sections were then washed 3x with PBS and fixed with 1.5 mM BS(PEG)9 in PBS for 20 minutes, inactivated in PBS + 100 mM Tris pH 8 for 5 minutes, washed 3x in PBS, and stored in tightly sealed tissue culture dishes at 4° in 70% ethanol for 24 hours to one month.

After antibody staining, the samples were then modified with MelphaX ^54^ that was diluted 1:10 in 20 mM MOPS pH7.7 at 37oC for 1 hr. The samples were washed three times with PBS, then embedded in an acrylamide gel (4% v/v 19:1 acrylamide:bis-acrylamide (Bio-Rad), 300 mM NaCl, 60 mM Tris pH 8, 0.2% v/v tetramethylethylenediamine [TEMED], and 0.2% w/v ammonium persulfate [APS]) for 1.5 hrs at room temperature. The samples were then digested at 42oC for 48 hours in digestion buffer (2% v/v sodium dodecyl sulfate [SDS] (Thermo Fisher), 1% v/v proteinase K (New England Biolabs), 50 mM Tris pH 8 (Ambion), 300 mM NaCl (Ambion), 0.25% Triton X-100 (Sigma), 0.5 mM ethylenediaminetetraacetic acid [EDTA] (Ambion)).

After digestion, samples were washed three times with PBS + 0.1% Triton X-100 to remove residual SDS. Samples were then hybridized with a mixture containing 2×SSC, 30% v/v formamide (Ambion), 1% v/v murine RNase inhibitor (New England Biolabs), 0.1% w/v yeast tRNA, 5% w/v PEG35000 (Sigma), 1 uM of a polyA probe, a library of probes for imaging pre-rRNA, mtRNA, and *Albumin* at 1 nM/probe, and each RCA-MERFISH encoding probe library at 1 nM/probe. The polyA probe had a mixture of DNA and LNA nucleotides (/5Acryd/TTGAGTGGATGGAGTGTAATT+TT+TT+TT+TT+TT+TT+TT+TT+TT+T) where T+ is a locked nucleic acid and /5Acryd/ is a 5’ acrydite modification. The samples were then hybridized for 36-48 hours at 37oC in a humidified chamber.

After hybridization, the samples were washed twice for 30 min in 2×SSC, 30% v/v formamide at 47oC. The samples were then washed three times with PBS + 0.1% v/v Tween 20, and once briefly with preligation buffer (50 mM Tris-HCl pH8, 10 mM MgCl2). The RCA-MERFISH probes were ligated with 10% v/v (∼1 uM) SplintR ligase (New England Biolabs), 1× SplintR ligase buffer, 1% v/v murine RNAse inhibitor, and 100 nM RCA primer (TCTTCACCCGGGGCAGCTGAA*G*T, where * is a phosphorothioate bond) at 37oC for 1 hr. Samples were then washed three times with PBS + 0.1% Tween 20. Next, samples underwent rolling circle amplification using 1× Phi29 Buffer (Lucigen), 10% v/v Phi29 enzyme (Lucigen), 0.2 mg/ml BSA, 250 µM dNTP (New England Biolabs), 25 µM aminoallyl-dUTP (ThermoFisher), and 1% v/v murine RNase inhibitor for 2 hrs at 37oC. Finally, the samples were washed three times in 1×PBS, and then crosslinked for 30 min with 1×PBS + 1 mM BS(PEG)9 at room temperature, before a final quick wash with 1×PBS. Samples were stored in 1×PBS with 1% v/v murine RNAse inhibitor at 4oC for up to one week.

#### Multiplexed RNA and protein imaging by RCA-MERFISH and sequential hybridization

RCA-MERFISH samples were imaged on a custom epifluorescent microscope with automated fluidics, as previously described for MERFISH imaging ^126^. Briefly, samples were mounted in a flow cell (Bioptechs) with a 0.75-mm-thick flow gasket on a Nikon epifluorescence microscope.

For each round of hybridization, mixtures of Cy7-, Cy5-, Cy3-labeled readout probes for each triplet of bits to be read out were diluted to a final concentration of 10 nM/probe in 5 mL of 2×SSC, 10% formamide, 0.1% Triton X-100. The samples were stained for 15 min, then washed with 2×SSC, 10% formamide, 0.1% Triton X-100. Finally, imaging buffer was flowed into the chamber. The imaging buffer consisted of 2×SSC, 10% w/v glucose (Sigma), 60 mM Tris-HCl pH8.0, ∼0.5 mg/mL glucose oxidase (Sigma), 0.05 mg/mL catalase (Sigma), 50 µM trolox quinone (generated by UV irradiation of 6-hydroxy-2,5,7,8-tetramethylchroman-2-carboxylic acid (Sigma)), 0.2% v/v murine RNAse inhibitor, and 0.1% v/v of Hoechst 33342 dye (ThermoFisher).

After the readouts were hybridized and imaging buffer added, the samples were imaged with a high-magnification, high-numerical aperture objective. For wildtype animals in Figs 1-2, a 60X 1.4NA oil immersion objective was used, with a pixel size of 108 nm/pixel. For genetically perturbed experiments in Figs 3-7, a 40X 1.3NA oil immersion was used, with a pixel size of 162 nm/pixel. We imaged each field of view (FOV) with a 10-plane z stack with 1.5 µm spacing between adjacent z planes, where each z-plane was imaged in the 750-nm, 650-nm, 560-nm, and 405-nm channels. After each round of imaging, the readout probes were stripped off using 2×SSC, 80% formamide stripping buffer for 10 min, followed by two washes of readout buffer, and one wash of 2×SSC. The different panels (antibodies and structural RNAs, endogenous RNA, and barcode RNA) were imaged back-to-back on the same tissue sections, where protein (labeled with oligo-conjugated antibodies) and structural RNA were first imaged by with sequential rounds of multicolor FISH, followed by endogenous RNAs and barcode RNAs in two separate RCA-MERFISH runs. Each experiment took 36-48 hours depending on the number of fields of view, and whether the perturbation barcode library was imaged in addition to the endogenous RNA and protein panels.

#### Data Processing Pipeline

RCA-MERFISH gene expression was processed using a modified version of the MERlin pipeline, as previously described ^126,128^, with the addition of a machine learning filtering step that used XGBoost to train a classifier to discriminate incorrectly decoded molecules that were assigned to blank barcodes and putatively correctly decoded molecules that were assigned to coding barcodes on a subset of the data. This classifier was then applied to the remainder of the data, and only molecules that were classified as coding molecules with an adaptive 5% false-positive threshold were exported for assignment to individual cells. Each library (endogenous RNA or barcodes) were decoded separately.

Cell segmentation was accomplished using a custom Cellpose model that was trained on images of polyA staining for cytoplasm + nucleus, and Na+/K+ ATPase staining for membrane. A separate model was trained for data collected using the 60X and 40X objectives, and applied to the respective datasets. Each cell was assigned a unique identifier and its position, shape, and z extent were recorded. The molecules that were exported from each field of view were then assigned to cells based on overlap between molecules and Cellpose-created mask for each cell in three dimensional space. The total number of molecules of each type for each cell was summed, and the result exported as an AnnData objective.

After decoding and cell assignment, all RCA-MERFISH datasets were concatenated into a single dataset. This was done separately for the wildtype physiological perturbation data (Figures 1 and 2) and the genetically perturbed data (Figures 3-7). Cells were filtered to remove all cells with less than 25 or greater than 1500 molecules per cell, and genes expressed in <3 cells were removed. Each cell’s gene expression values were scaled such that the sum of gene expression values per cell added to 10,000, then was log-transformed. The area and number of molecules per cell were regressed out using linear regression, and the residuals were z-scored. The first 20 principal components were then computed and the data were integrated with log-transformed, z-scored Flex data using Harmony, using the top 40 principal components computed based on the subset of genes measured in RCA-MERFISH. The data were then jointly clustered using Leiden clustering with resolution = 0.4. Clusters with fewer than 10 differentially expressed genes were then greedily merged, and the final set of clusters were manually annotated based on marker gene expression. For perturbation data, the individual guide was called for each cell when there were > 3 molecules per cell for a given barcode.

The labels assigned to clusters in the integrated Flex and RCA-MERFISH data from wildtype animals were transferred to the perturbed data through integration and label transfer. After pre-processing to filter out cells, normalize, log transform, and z-score the data as described above, the two datasets were integrated using Harmony. A K-nearest neighbors classifier was then trained to predict the clusters annotations of the wildtype RCA-MERFISH data from the top 20 principal components of the data, with K=10. This classifier was then applied to the perturbed RCA-MERFISH data to predict cluster identity.

For morphological imaging, each channel used for morphological imaging was adaptively contrast adjusted. A single z-plane in the middle of each segmented cell was selected, and the image of each cell in each channel was cropped out of the larger field of view, in a 256 x 256 square, using the segmentation for that cell as a mask. The background around each cell outside of the mask within that 256 x 256 square was set to zero. The imaging data for each cell was then associated with the unique identifier for that cell, for integration with RCA-MERFISH data for endogenous RNA and perturbation barcodes.

#### Deep autoencoder model

We developed a deep autoencoder model to extract biologically relevant features from single-cell multiplexed protein and abundant RNA images based on the second generation of the vector quantized variational autoencoder (VQ-VAE) ^64^. The model takes advantage of the power of VQ-VAE to learn meaningful representations using self-supervised training. Inspired by the cytoself model ^13^, we used auxiliary classification tasks to guide the model to focus on features of biological importance. Each single-color protein image of a particular protein/RNA channel was concatenated to a fiducial channel to create a two-color image. We used the polyA FISH staining as the fiducial channel to provide information of the relative location of the stained proteins/RNAs to the nuclei. From the two-color image, the model used a ResNet with two residual blocks to generate a bottom level representation as described in the second generation VQ-VAE design. The bottom level representation was then used to generate a top level representation with another ResNet with two residual blocks. Both bottom and top representations were converted to discrete representations by vector quantization ^64^. The quantized bottom and top representations were concatenated and decoded by a ResNet with two residual blocks to regenerate the input image. The concatenated representation vectors were fed into a multi-layer perceptron (MLP) classifier with one hidden layer to predict the identity of the protein/RNA channels. The hidden representation of the MLP classifier had 512 dimensions, which was used as the representation vector of the input image. The representation vectors for different protein/RNA channels of one cell were concatenated to generate the morphological representation of a cell. The cell representation was fed into another MLP classifier to predict the transcriptionally defined cell type and/or the diet condition of the cell.

The loss function of the autoencoder model contained 4 terms, namely, the latent loss, the image reconstruction loss, the protein/RNA classification loss and the cell-type or diet-condition classification loss. We used the same latent loss definition as the VQ-VAE model ^64^, which measured the difference between the latent representations before and after quantization. The image reconstruction loss was defined as the mean squared error of the reconstructed image compared to the input image. The protein/RNA and cell-type/diet-condition classification losses were defined as the cross entropy loss of classification.

#### Training the deep autoencoder

Images of single-cells were cropped out from the fields of view as square boxes. We set all the pixels outside the target cells to zero using the cell segmentation masks, such that each cropped image only contained one cell. The cropped single-cell images were rescaled to 128x128 pixels and used as input for the autoencoder training. We optimized the autoencoder model with stochastic gradient descent using the Adam optimizer until the loss function converged.

We trained a VQ-VAE model to embed cells under different diet conditions using both the protein classification auxiliary task and the cell classification auxiliary task that predicted transcriptionally defined cell types and diet conditions. For embedding the cells under CRISPR perturbations, because we only included hepatocytes, the autoencoder model was trained using only the protein classification auxiliary task.

#### Tissue zone segmentation

The zonal segmentation of tissue in Fig 6E-G was accomplished by computing for each replicate a 2D histograms at 50 µm resolution of the number of Hep1+Hep2 or Hep5+Hep6 cells in each bin. These histograms were then blurred with a Gaussian filter with sigma = 0.5 and normalized by dividing by the maximum across all bins. For each bin, the zone was determined as whether the normalized Hep1+Hep2 or Hep5+Hep6 count was greater for that bin.

### QUANTIFICATION AND STATISTICAL ANALYSIS

#### scRNA-seq Analyses

##### Energy Distance Calculation and Permutation Testing

Energy distances and permutation tests were calculated according to the method of ^71^. We used the first 20 PCs generated from the tp10k-normalized, log1p-transformed data. We used the Holm-Šídák multiple testing correction for our permutation testing. For the fasted vs *ad libitum* comparisons, we compared the transcriptional states of perturbed cells to those of cells with control sgRNAs in the same mouse in the same condition.

##### Z-scoring relative to control

For some purposes, we analyzed z-scored transcriptional data. We calculated the z-scores by tp10k-normalizing the data, identifying the mean and standard deviation for each mRNA in control cells, and then used these values to identify the z-score for each gene in each cell, relative to control cells. We performed this calculation separately for each barcode in each GEM group and then concatenated the cells to form a final z-normalized expression matrix. This transformation emphasizes changes in mRNAs with low variance in control cells and may decrease batch effects between barcodes and GEM groups.

##### Pseudobulk Correlation Calculations

Correlations calculated between pseudobulk transcriptional responses are either Pearson correlations calculated from mean log-transformed transcriptional responses, with the expression in controls subtracted to a Perturbation-associated phenotype, or correlations of mean Z-scored transcriptional changes. We did not observe significant differences between the two. To decrease noise, we only included genes with high expression in control cells (highest 250 or highest 1000) in the calculations. In Figure 4M, we show sgRNAs targeting genes that have two sgRNAs that have substantial transcriptional phenotypes.

##### Differential Expression

We used Benjamini-Hochberg-corrected Mann-Whitney testing to identify genes whose expression is significantly affected by perturbations (corrected p < 0.05). We used tp10k-normalized, log1p-transformed data for these calculations. The heat map showing changes in expression (Figure 7D) represents log2-fold changes of pseudobulk expression versus cells with control perturbations.

##### Scoring and significance testing

The scores in Figures 5A and 5B are calculated using gene sets derived from a previous large-scale perturbation experiment in cell culture ^18^. The scores are calculated as the pseudobulk z-scored change relative to control cells, averaged for all genes in the gene set, for all cells with each perturbation. The scores in Figures 6A, 6B, and 7C are calculated with a literature-curated set of zonation markers and reflect the sum of pseudobulk or single cell z-scored changes relative to control cells. The periportal markers are *Cyp2f2*, *Hal*, *Hsd17b13*, *Sds*, *Ctsc*, *Aldh1b1*, and *Pck1* and the pericentral markers are *Cyp4a14*, *Cyp2d9*, *Gstm3*, *Cyp4a10*, *Mup17*, *Slc1a2*, *Slc22a1*, *Cyp1a2*, *Aldh1a1*, *Cyp2a5*, *Gulo*, *Cyp2c37*, *Lect2*, *Cyp2e1*, *Oat*, *Glul*. Periportal expression contributes positively to the score and pericentral contributes negatively to the score, or alternatively they are shown separately. Significance is calculated relative to negative control cells by Mann-Whitney with Benjamini-Yekutieli correction for multiple testing (corrected p < 0.05).

##### Clustering and genotype-phenotype mapping

The perturbation heatmap in 4D is generated from the hierarchical clustering of a joint vector including Pearson correlations of pseudobulk log-transformed transcriptional responses measured by sequencing and pseudobulk z-scored staining intensity changes measured by imaging (Euclidean distance metric, UPGMA algorithm). The representation of gene co-regulation in Figure 4E is generated from correlations between z-scored pseudobulk expression levels of pairs of genes across perturbation (the transpose of the transcriptional component of the data used to generate the correlations in Figure 4D). The position of spots derives from a two-dimensional minimal distortion embedding of these correlations that tries to place co-regulated genes in proximity. The color of spots derives from a density-based clustering of a separate twenty-dimensional minimal distortion embedding of co-expression, calculated according to the method used for mRNAs in Figure 4B of ^18^.

#### Intensity Analysis

##### Z-scoring relative to control

When analyzing intensity, we z-scored each protein/RNA channel relative to the mean and standard deviation of that channel in control cells (cells with control sgRNAs), from that same imaging sample. We then concatenated the cells from the various imaging samples to form a final z-normalized intensity matrix. This transformation emphasizes changes in protein/RNA intensities with low variance in control cells and may decrease batch effects between imaging samples.

##### Pseudobulk Correlation Calculations

Correlations calculated between pseudobulk protein/RNA intensity responses are Pearson correlations calculated from mean z-scored intensity responses. In 3Q, we only show perturbations targeting genes that have two sgRNAs that are significant in a corrected energy distance permutation test.

##### Number of Protein/RNA Channels Exhibiting Differentially Intense Signals

To quantify the number of differentially intense protein/RNA channels (Figure S9F), we used Benjamini-Hochberg-corrected Mann-Whitney testing (corrected p < 0.05) on the z-scored intensity data.

##### Intensity Scoring and significance testing

The scores in Figures 5C, 5D, 5E, 5F, 7A, 7B, and 7G are calculated as the mean z-scored change relative control cells for all cells with each perturbation. Significance is calculated relative to negative control cells by Mann-Whitney with Benjamini-Yekutieli correction for multiple testing, corrected p < 0.05.

### KEY RESOURCES TABLE

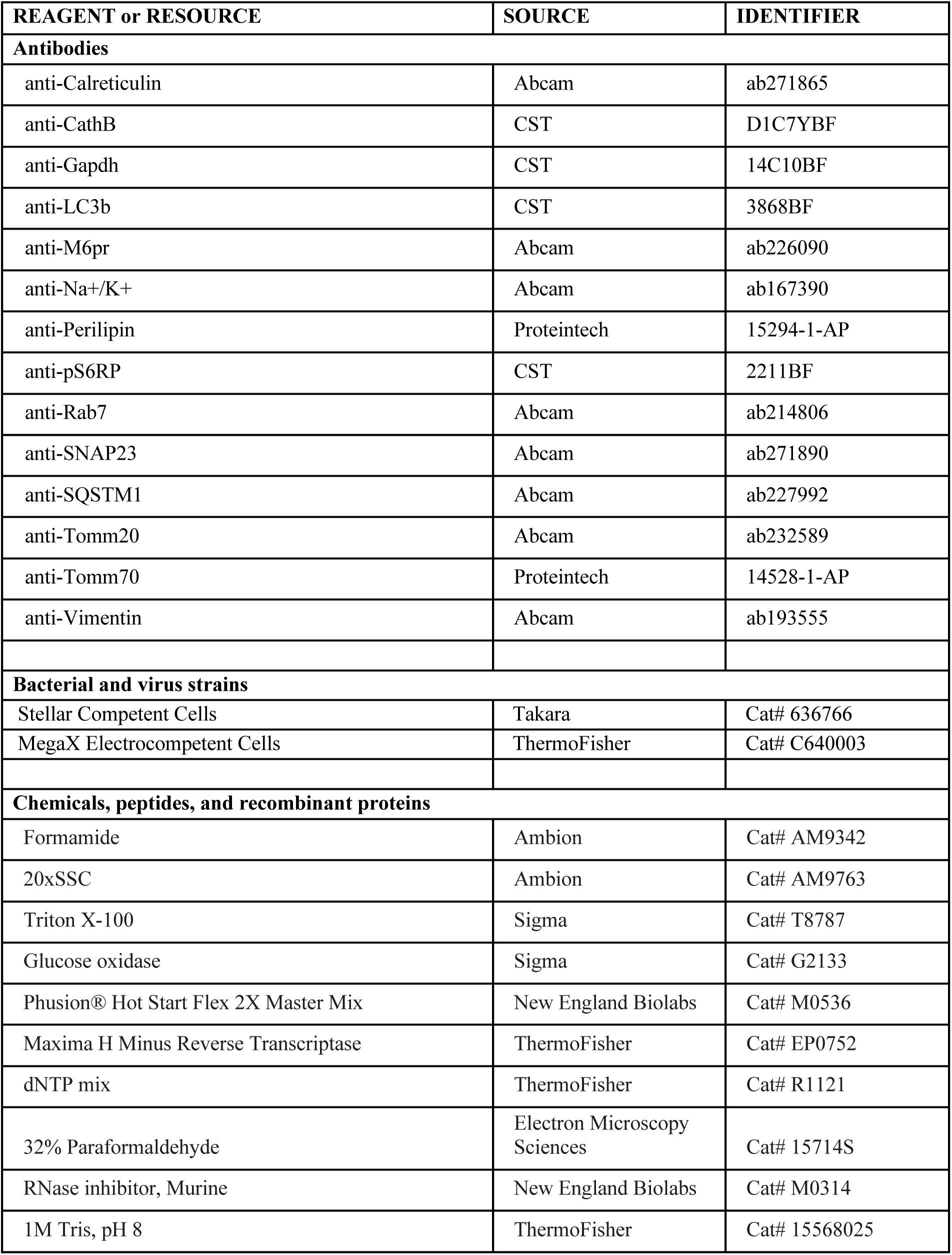
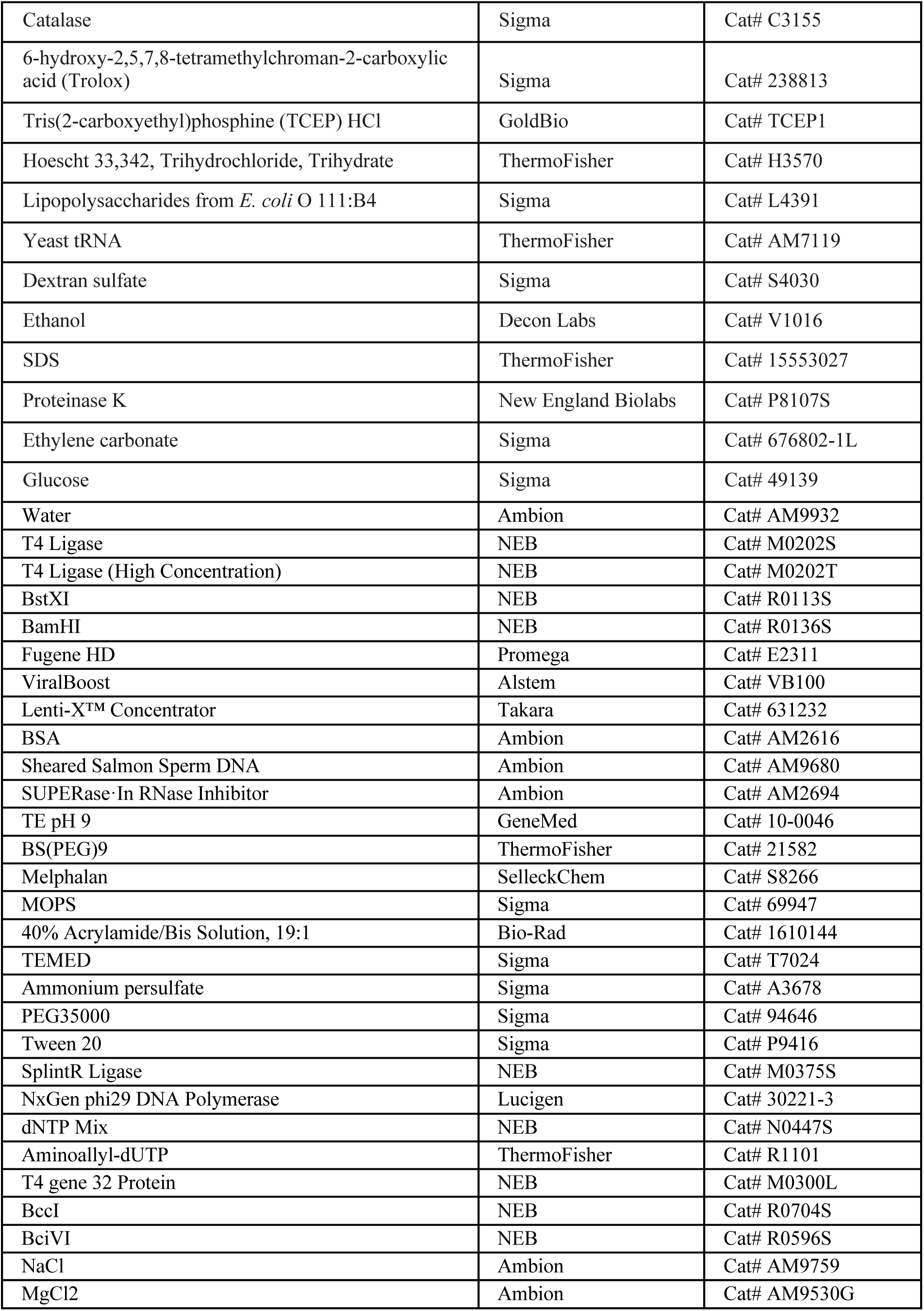
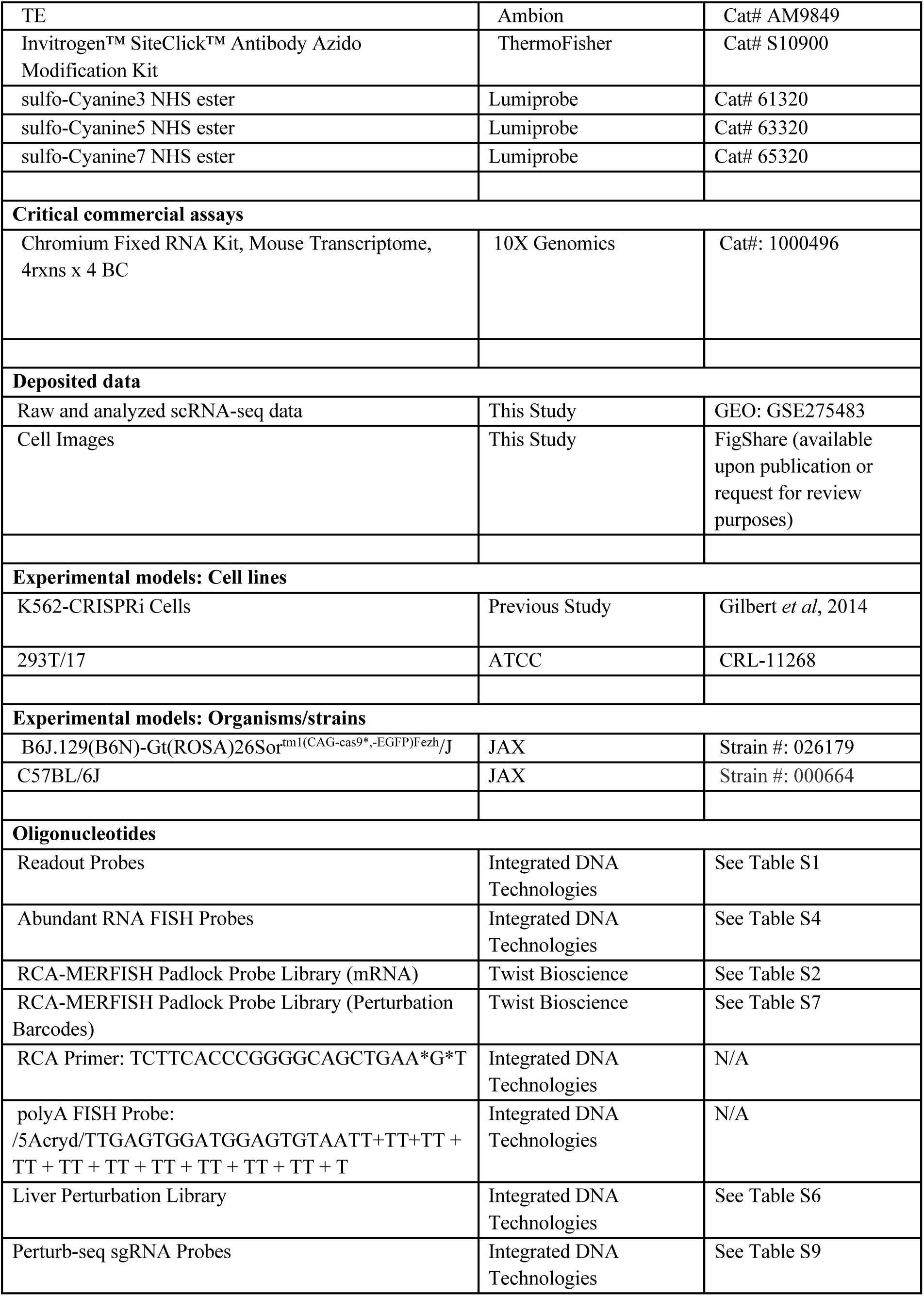
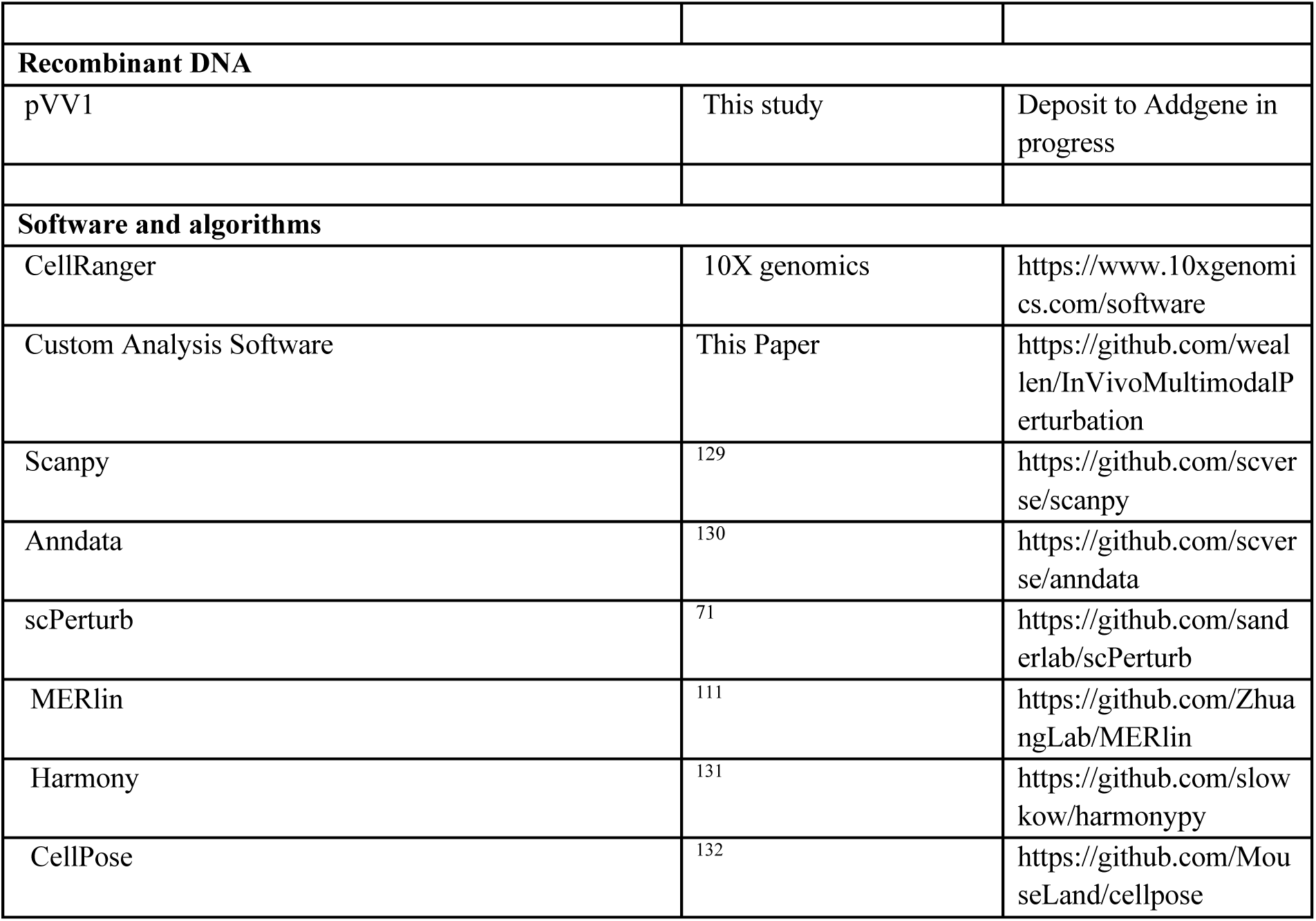

---

**Figure S1:**
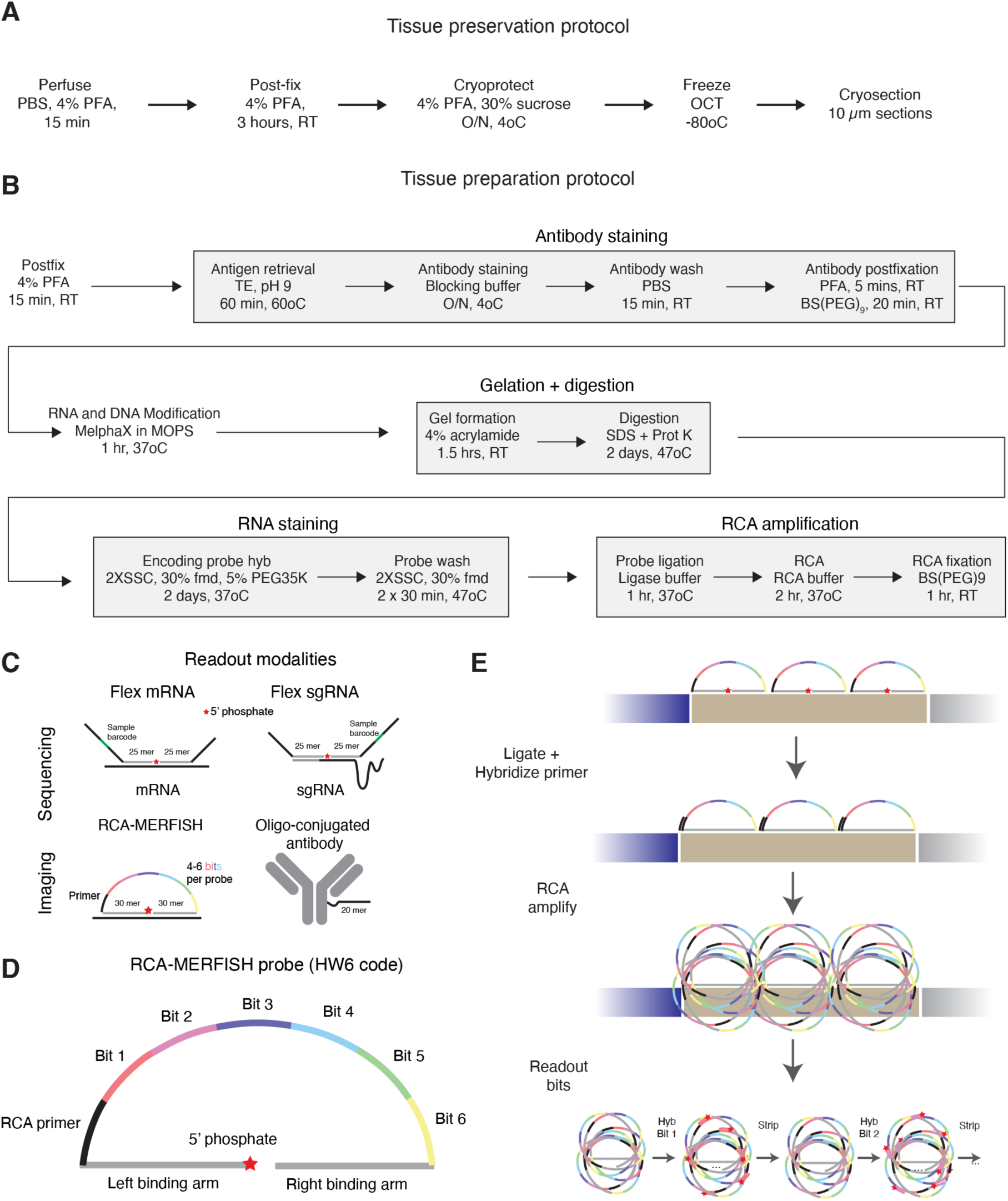
Workflow and optimizations for imaging-based screening A. Experimental procedure for tissue preservation by PFA fixation and cryoprotection. B. Detailed experimental protocol for multimodal oligo-conjugated antibody staining and RCA-MERFISH in fixed tissue. C. Different readout modalities by imaging or sequencing D. Diagram of padlock probe design for RCA-MERFISH with a Hamming Weight (HW) 6 code. The readout sequences (marked as bit 1 to bit 6), the presence of which determined the MERFISH code, are directly encoded in the padlock probe. E. Diagram of RCA-MERFISH signal amplification process. Probes that have both arms hybridized to target RNA and adjacent to each other are ligated, and then the ligated probes are amplified through rolling circle amplification (RCA). After RCA, the individual bits in each amplicon are read out through fluorescent microscopy, over multiple rounds of staining with fluorescent readout probes, and then dehybridizing (‘stripping’) the probes off with a high formamide concentration wash.

**Figure S2:**
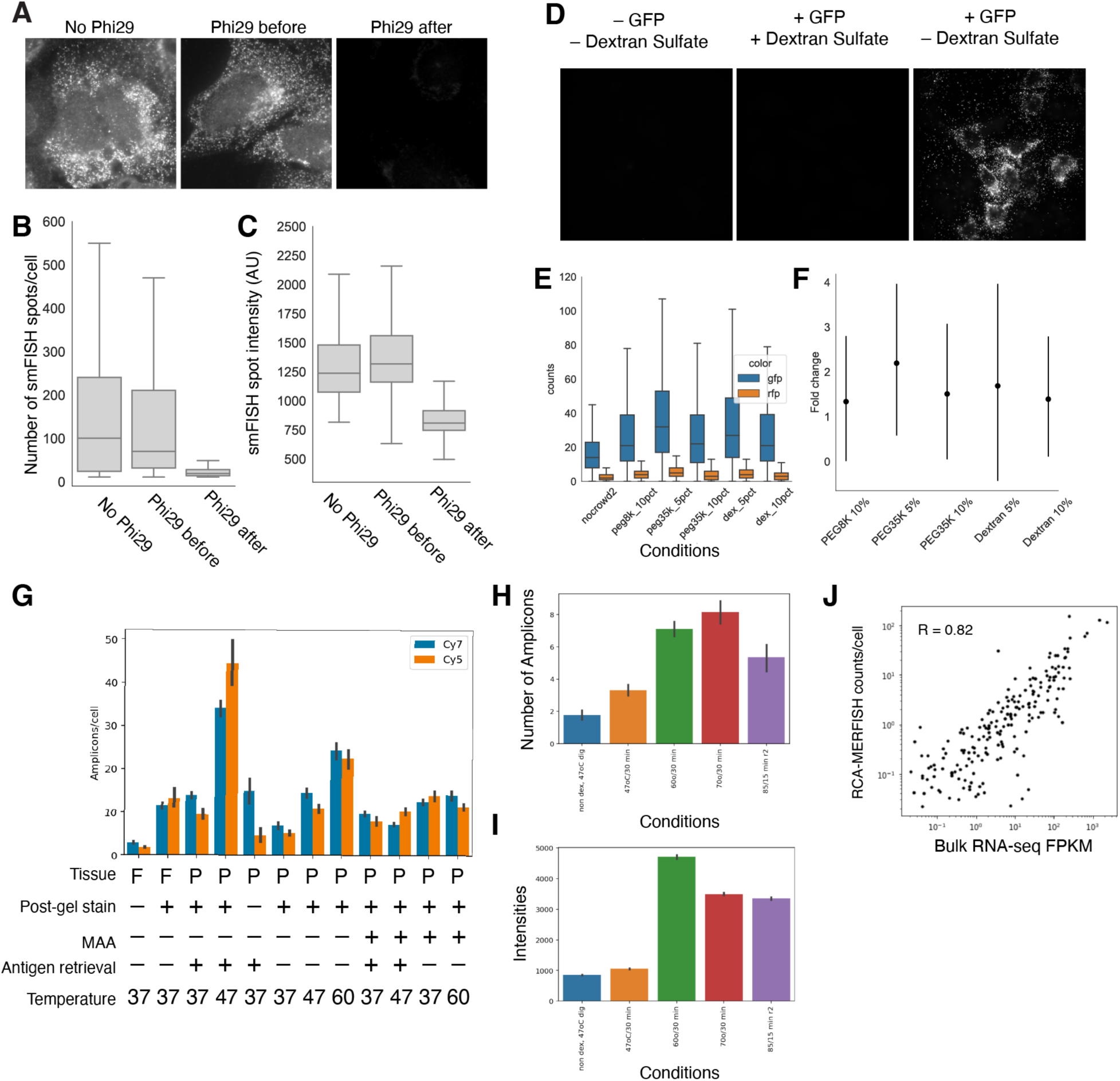
Optimization of RCA-MERFISH protocol A. Phi29 used in RCA degrades ssDNA smFISH probes. Left: smFISH signal in U-2 OS cells without Phi29 treatment at 37oC for 1 hr. Center: smFISH signal in U-2 OS cells pre-treated with Phi29 at 37oC for 1 hr before FISH staining. Right: smFISH signal in U-2 OS cells treated with Phi29 at 37oC for 1 hr after FISH staining. B. Quantification of effect of Phi29 on ssDNA smFISH probes from (A), in spots/cell. C. Quantification of effect of Phi29 on ssDNA smFISH probes from (A), in intensity/spot. D. Dextran sulfate inclusion in hybridization buffer inhibits Phi29 enzymatic activity. Left: amplified RCA-MERFISH padlock probe against GFP in U-2 OS cells not expressing GFP, detected by readout probes complementary to readout sequences on the padlock. Center: amplified RCA-MERFISH probe against GFP in U-2 OS cells expressing GFP, with dextran sulfate in the hybridization buffer, detected by readout probes complementary to readout sequences on the padlock. Right: amplified RCA-MERFISH probe again GFP in U-2 OS cells expressing GFP, without dextran sulfate in the hybridization buffer, detected by readout probes complementary to readout sequences on the padlock. E. Optimization of alternative crowding agents to dextran sulfate. Multiple additives to hybridization mixture, staining U-2 OS cells expressing either GFP or mCherry with a single probe against GFP, in terms of number of spots per cell, distinguishing GFP+ (signal) and mCherry+ (background) cells. Peg8k = Poly(ethylene glycol) average mol wt 8,000, Peg35k = Poly(ethylene glycol) average mol wt 35,000, Dex = unsulfonated dextran; all are added to the hybridization solution so the final w/v is at the indicated percent. F. Quantification of increase in efficiency of different additives to hybridization mix from (E), relative to control (no additive to hybridization mix). G. Optimization of RCA-MERFISH. Fresh-frozen (F) and PFA-fixed (P) tissue was tested with staining RNA after gel embedding (+ Post-gel stain) or before gel embedding (– Post-gel stain), with or without the addition of methacrylic acid NHS ester (MAA) along with MelphaX, with or without antigen retrieval, and varying the digestion temperature from 37oC to 60oC. Counts are number of amplicons per cell across the first two bits, detected in the Cy7 and Cy5 color channel, from a 120 gene RCA-MERFISH library. H. Optimization of RCA-MERFISH across different decrosslinking conditions, quantified by amplicon counts per cell for a single bit, using an RCA-MERFISH library at low concentration (∼0.1 nM/probe), hence the lower counts per cell than (G) which used ∼10X higher probe concentration. The conditions tested are (1) no decrosslinking, (2) decrosslinking at 47° for 30 minutes, (3) decrosslinking at 60° for 30 minutes, (4) decrosslinking at 70° for 30 minutes, and (5) decrosslinking at 85° for 15 minutes. All decrosslinking was conducted in TE pH 9. I. Optimization of immunofluorescence across different decrosslinking conditions, measuring total intensity per field of view for a Tomm20 antibody. The decrosslinking conditions were the same as in Figure S2H. J. Correlation of 209 gene RCA-MERFISH (after optimization) with bulk RNA-seq from the liver.

**Figure S3:**
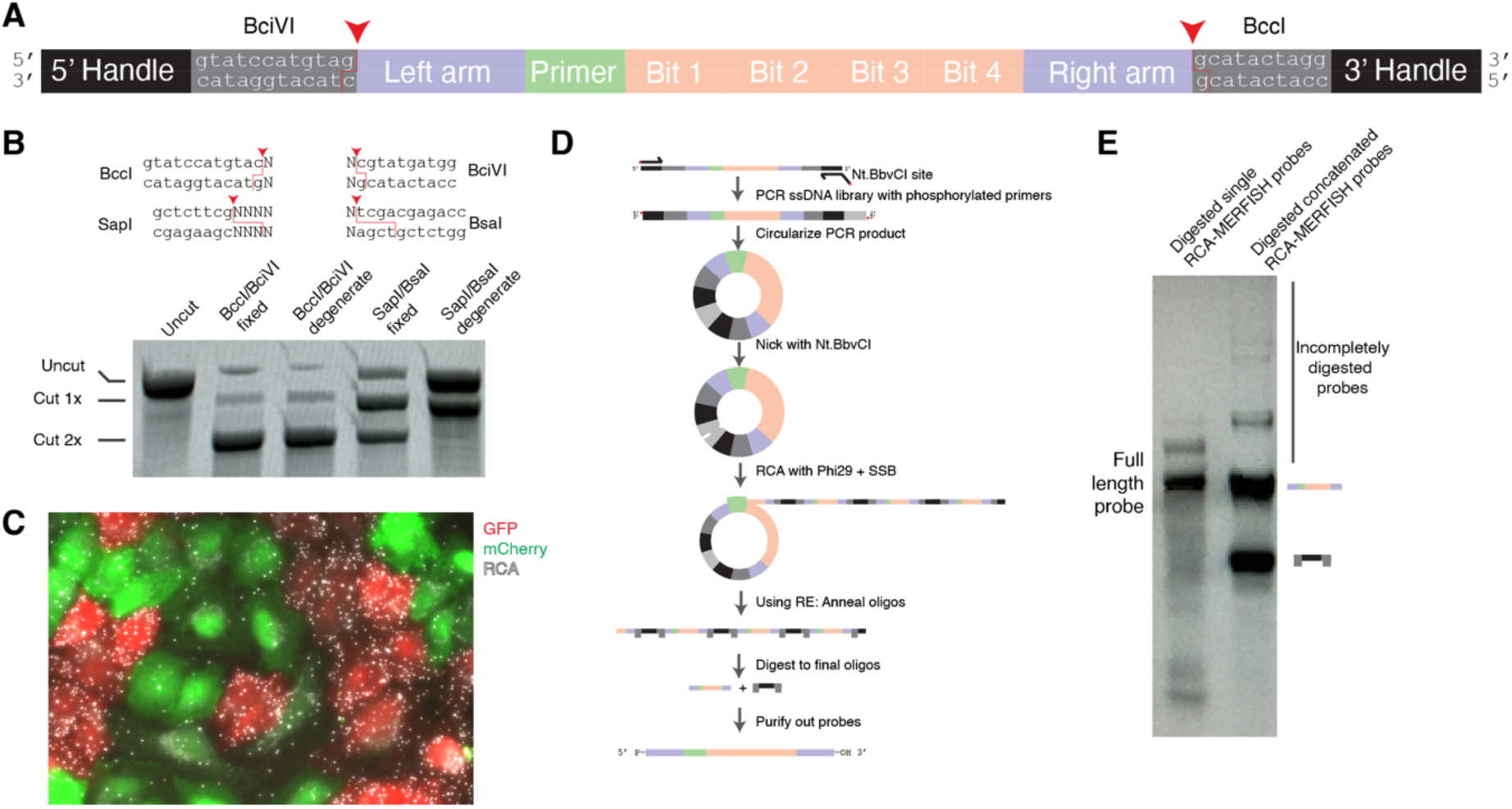
RCA-MERFISH padlock probe production A. RCA-MERFISH padlock probe design for pooled oligonucleotide synthesis, prior to amplification and probe library synthesis. B. Testing different Type IIS restriction enzymes on single restriction digested, PAGE-purified probe against GFP made through phophoramidite synthesis. C. RCA amplicons of digested RCA-MERFISH probe against GFP in U-2 OS cells expressing either GFP or mCherry. There are many more GFP amplicons in GFP-expressing cells than there are in mCherry-expressing cells, indicating the specificity of RCA-MERFISH. D. Probe library preparation protocol using RCA followed by restriction digestion. Femtomolar pools of oligos synthesized in arrays are amplified by PCR with phosphorylated primers containing an Nt.BbvCI site. The PCR amplicons are then circularized and nicked with Nt.BbvCI. The nick site is used to initiate rolling circle amplification, and the RCA product is then digested with BccI and BciVI after annealing on primers. E. Comparison of digested single probes and concatenated probes produced through RCA synthesis, on a 4% agarose gel electrophoresis.

**Figure S4:**
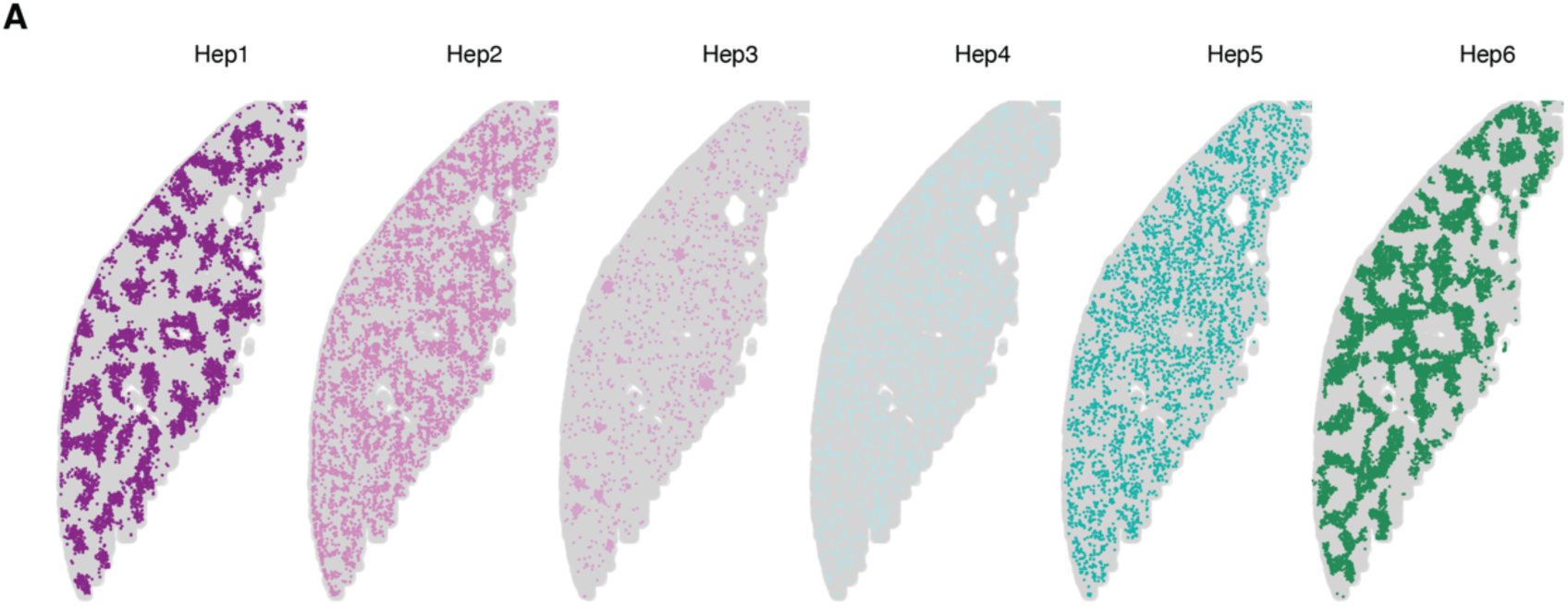
Hepatocyte transcriptional subtypes A. Spatial organization of different transcriptionally-defined hepatocyte subtypes, Hep1-6.

**Figure S5:**
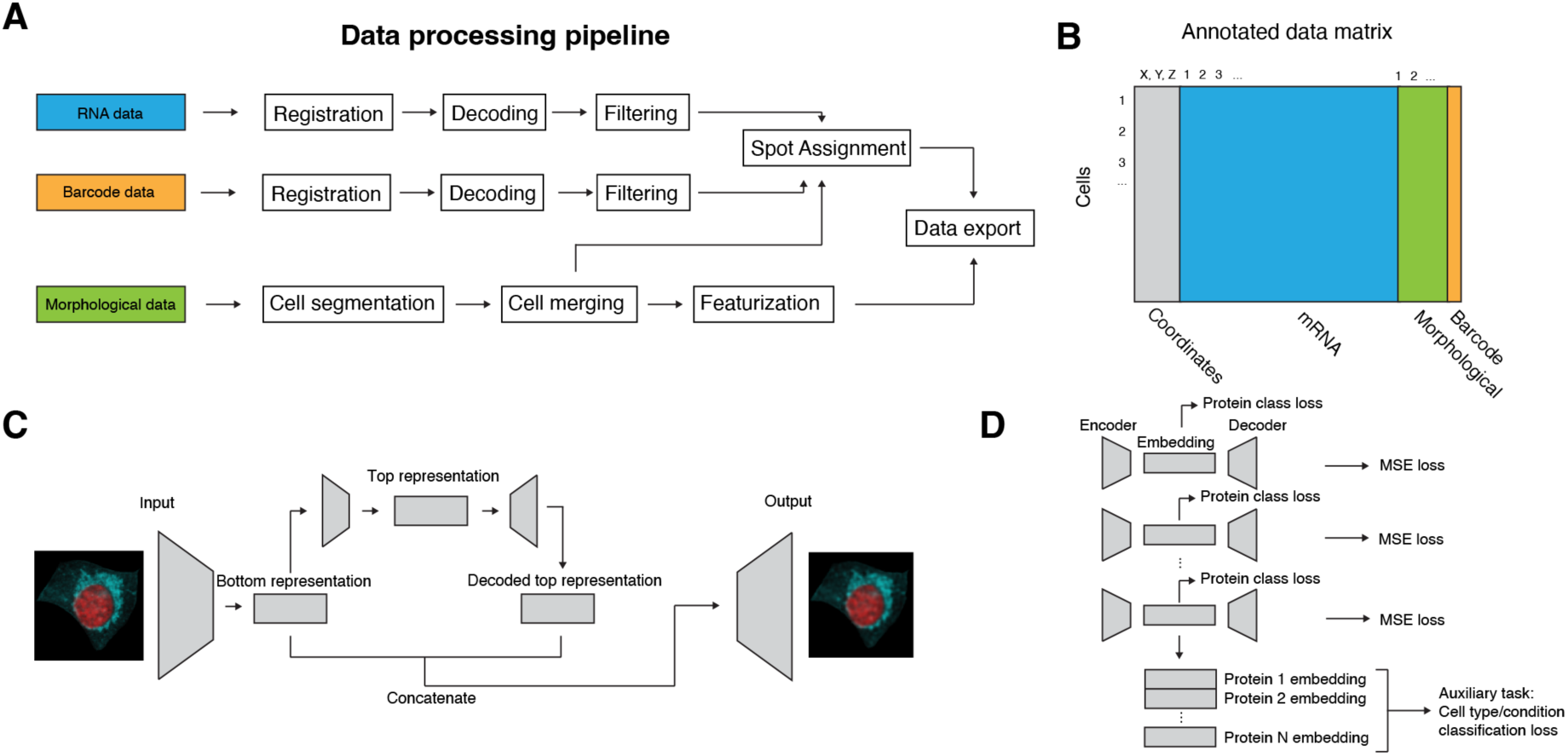
Data processing pipeline and deep learning model architecture A. Diagram of data processing pipeline. Each panel of barcodes, endogenous RNA or morphological data (protein or RNA) that is measured is collected back-to-back in the same experiment and then processed in parallel. The RCA-MERFISH data is processed by first registering to common fiducials across multiple rounds, then decoding the identity of individual molecules. The molecules are then filtered using machine learning on features of molecules (mean intensity, size, variance, difference between mean on- and off-bit intensity), to obtain a final 5% false positive rate that decode to a blank barcode. In parallel, the polyA and Na+/K+ ATPase channels of the morphological data are used to segment cells, which are then merged to eliminate duplicates of the same cells segmented in multiple fields of view. The cell segmentations are used to assign molecules to individual cells for quantification, and then the morphological channels are used with the segmentation mask to export the final images and per-gene quantification of expression for each cell. B. Diagram of final annotated data matrix combining all features. C. High-level diagram of VQ-VAE network across all channels. An input image is put into an artificial neural network that attempts to reconstruct the same image after passing the image through a low-dimensional bottleneck. In this case, two separate representations are created (top- and bottom-level) that attempt to capture different scales of features in the image. The bottom representation is formed first through one network, then further compressed to form a top representation with a separate autoencoder. The two representations are then concatenated and passed through a final network to reconstruct the original image. D. Detail of VQ-VAE network for each individual channel, trained simultaneously. A separate embedding is created for each morphological channel at the same time, using a mean squared error (MSE) loss to determine the accuracy of reconstruction. For each morphological channel, the embedding is used in a classification task to predict the identity of the protein or RNA being represented. The embeddings for each morphological channel are then concatenated and used to predict higher-level information about the type or state of each cell. When added to the overall training loss, these auxiliary predictive tasks are intended to constrain the representations that are formed by the VQ-VAE networks to capture salient features that discriminate different morphological channels and cell types or states.

**Figure S6:**
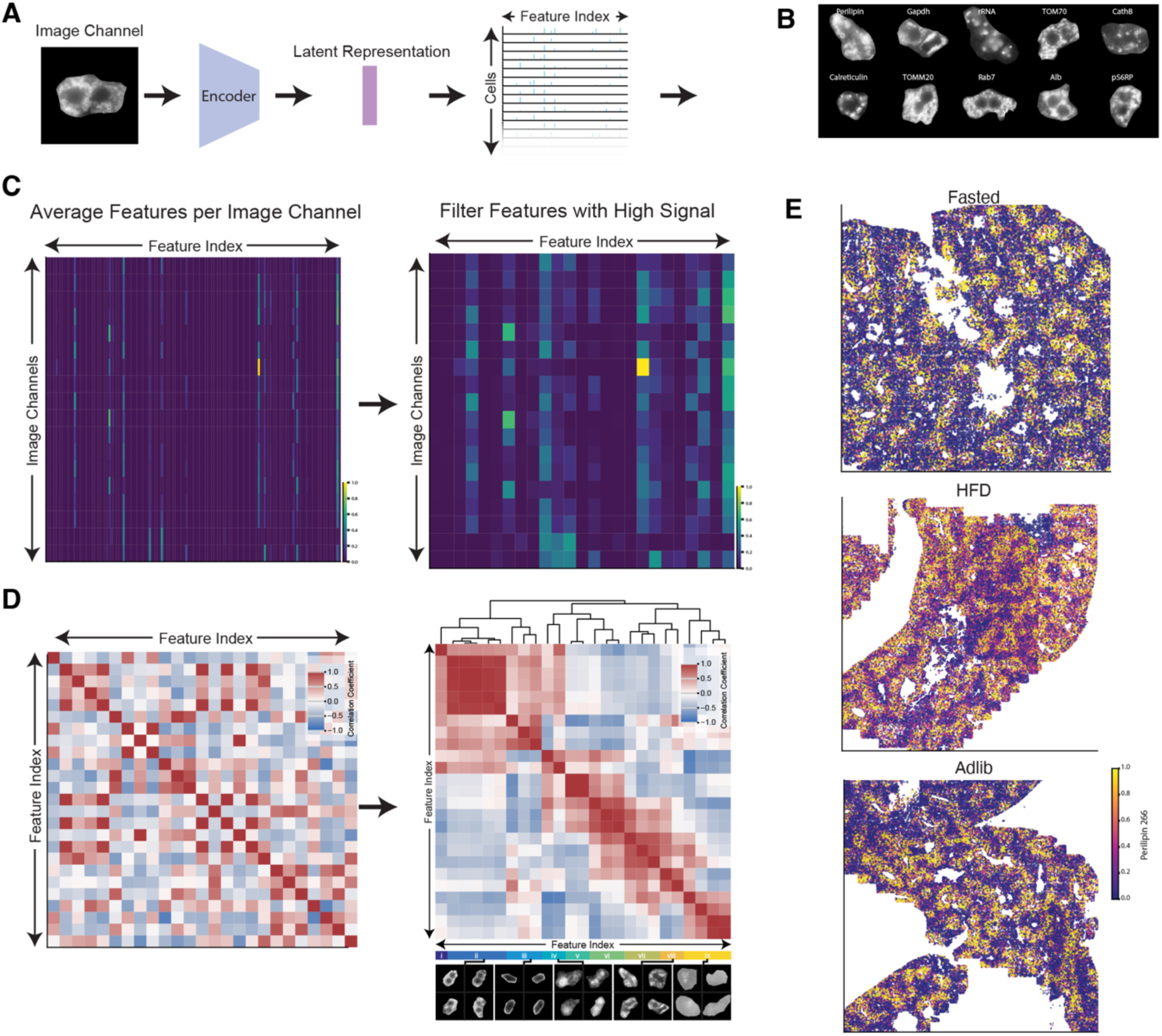
Analysis of image features from deep learning embedding A. Diagram of transformation of individual imaging channels into cell by feature representations. Each image channel for each cell is reduced to 512-dimensional vector. Here, we consider each dimension a feature. B. Examples of protein channel images that have high values for feature 266 (locally concentrated expression) from Class viii (as described below in (D)). This shows that the same protein/RNA feature measures the similar spatial patterns across different protein/RNA channels and cells. C. Heatmap of average feature weights across different imaging channels, with all features shown (left) or only features with high weight scores (high signals) shown (right). D. (Left) Heatmap of the pairwise correlation between high-signal features across cells. (Right) This heatmap is reordered through hierarchical clustering to reveal features that correlate strongly. Nine classes of features are manually identified and visualized. Each class of features captures similar spatial patterns. Cells with high weight scores of features from each class are displayed, including example classes (ii) cells with two Nuclei, (iii) signal enriched at cell membrane, (iv) signals showing relatively diffuse expression, (viii) signals showing locally concentrated expression, and (ix) noise. E. The spatial distribution of feature 266 in the Perilipin protein channel is illustrated for samples under fasted, high fat diet (HFD), and *ad lib* conditions. Cells displaying high Perilipin 266 values contain high amounts of concentrated perilipin clusters. Notably, pre-normalized values for the Perilipin 266 feature are considerably higher in the HFD sample.

**Figure S7:**
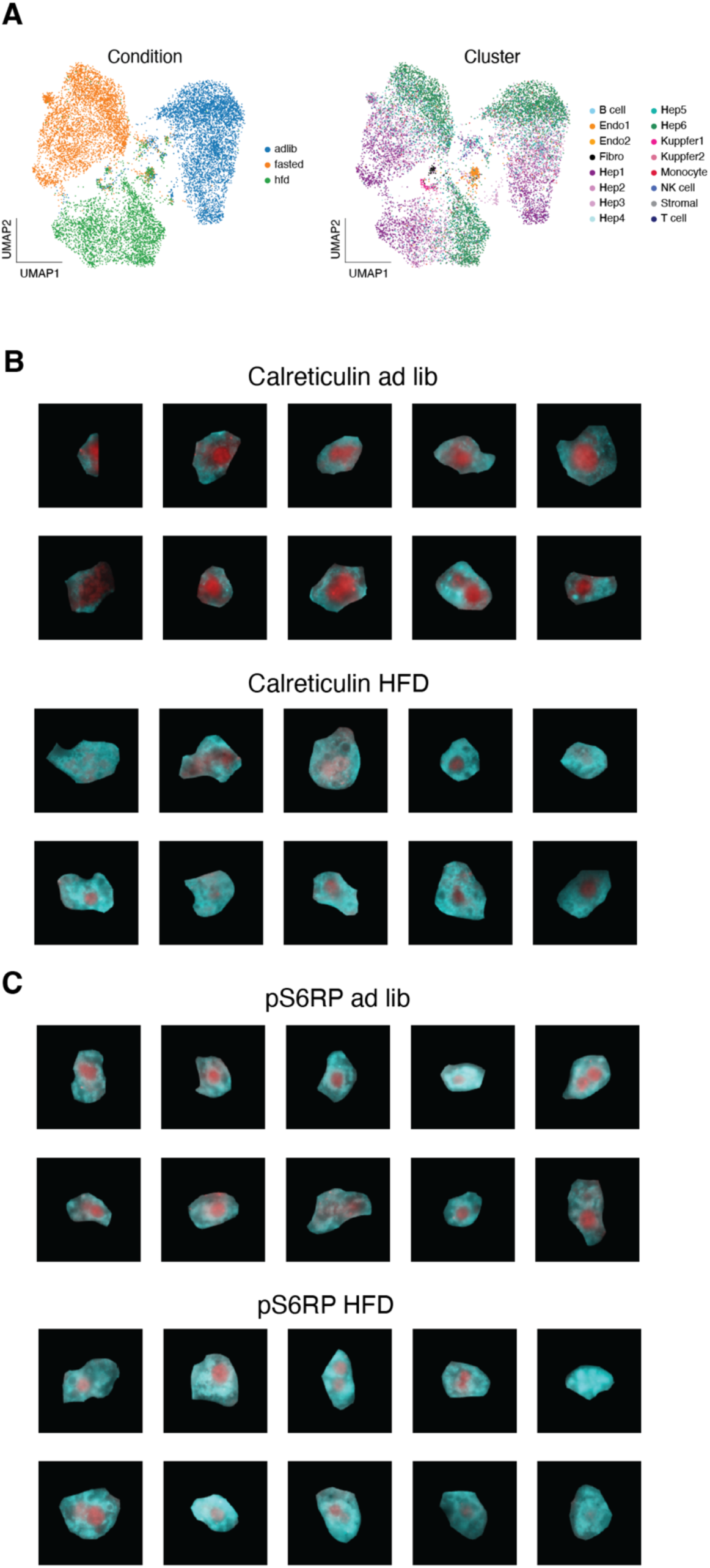
Changes in gene expression and morphology with physiological state A. UMAP of individual cells measured by 10X Flex from mice either with *ad lib* diet, overnight fasting, or 1 month high fat diet (HFD), colored by condition (left) or cell-type and subtype identity (right). B. Examples of calreticulin morphology in cells under *ad lib* or HFD conditions. C. Examples of pS6RP morphology in cells under *ad lib* or HFD conditions.

**Figure S8:**
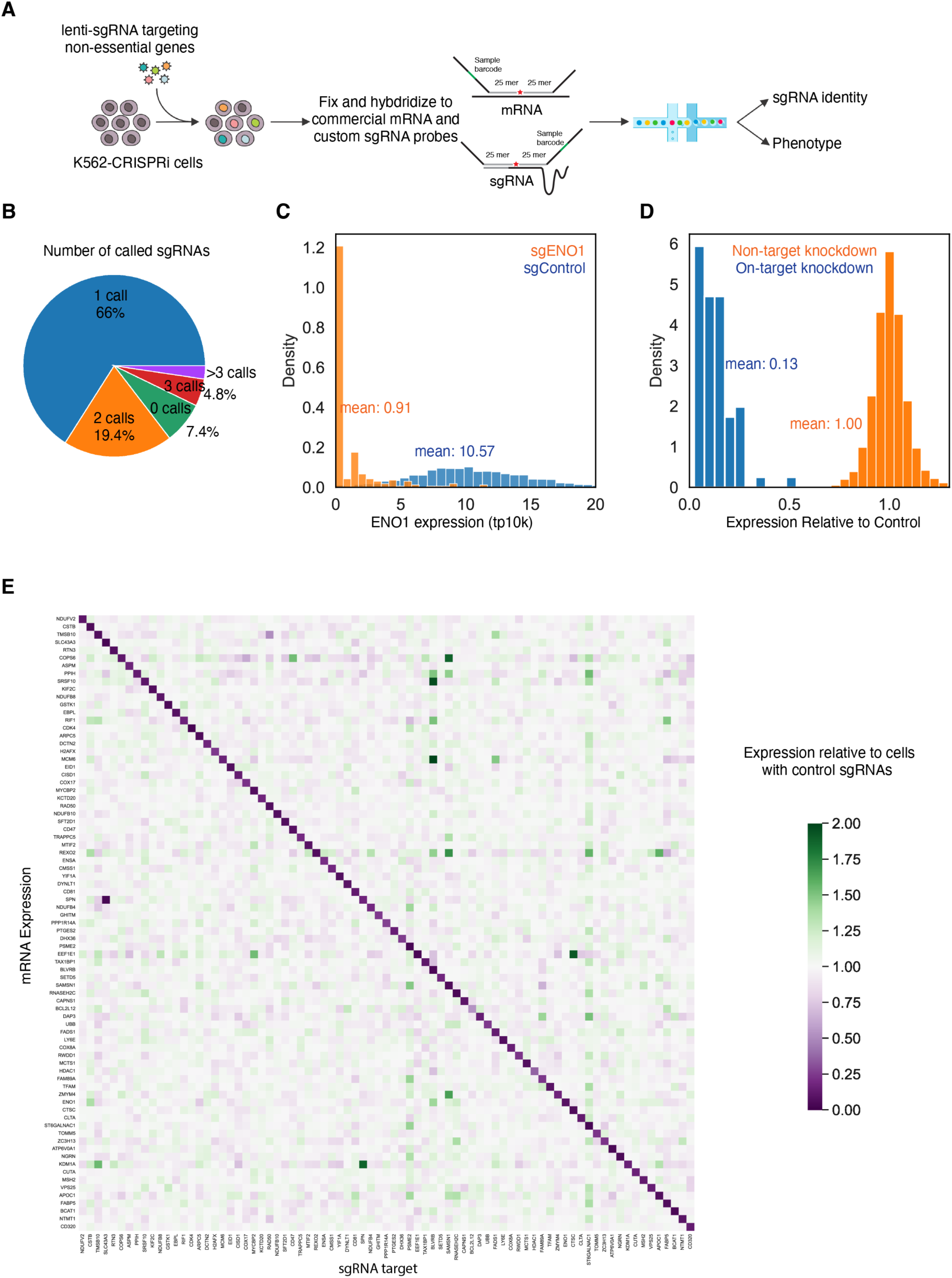
Validation of fixed cell Perturb-seq with CRISPRi A. Diagram of K562-CRISPRi Perturb-seq validation experiment B. Pie chart of the number of called sgRNA per cell. C. Histogram representing *ENO1* expression in cells with an sgRNA targeting *ENO1* or in cells with control sgRNAs. D. Histogram representing on-target knockdown and off-target knockdown, averaged across all targets in the experiment. On-target knockdown is defined as the expression of the target gene in cells with each corresponding sgRNA, relative to cells with control sgRNAs. Off-target knockdown is defined as the expression of each of other genes targeted in the experiment (not targeted in that cell), relative to expression of those genes in cells with control sgRNAs. E. Heat map representation of average expression of each of the indicated genes in cells with each of the indicated sgRNAs, relative to expression of those genes in cells with control sgRNAs.

**Figure S9:**
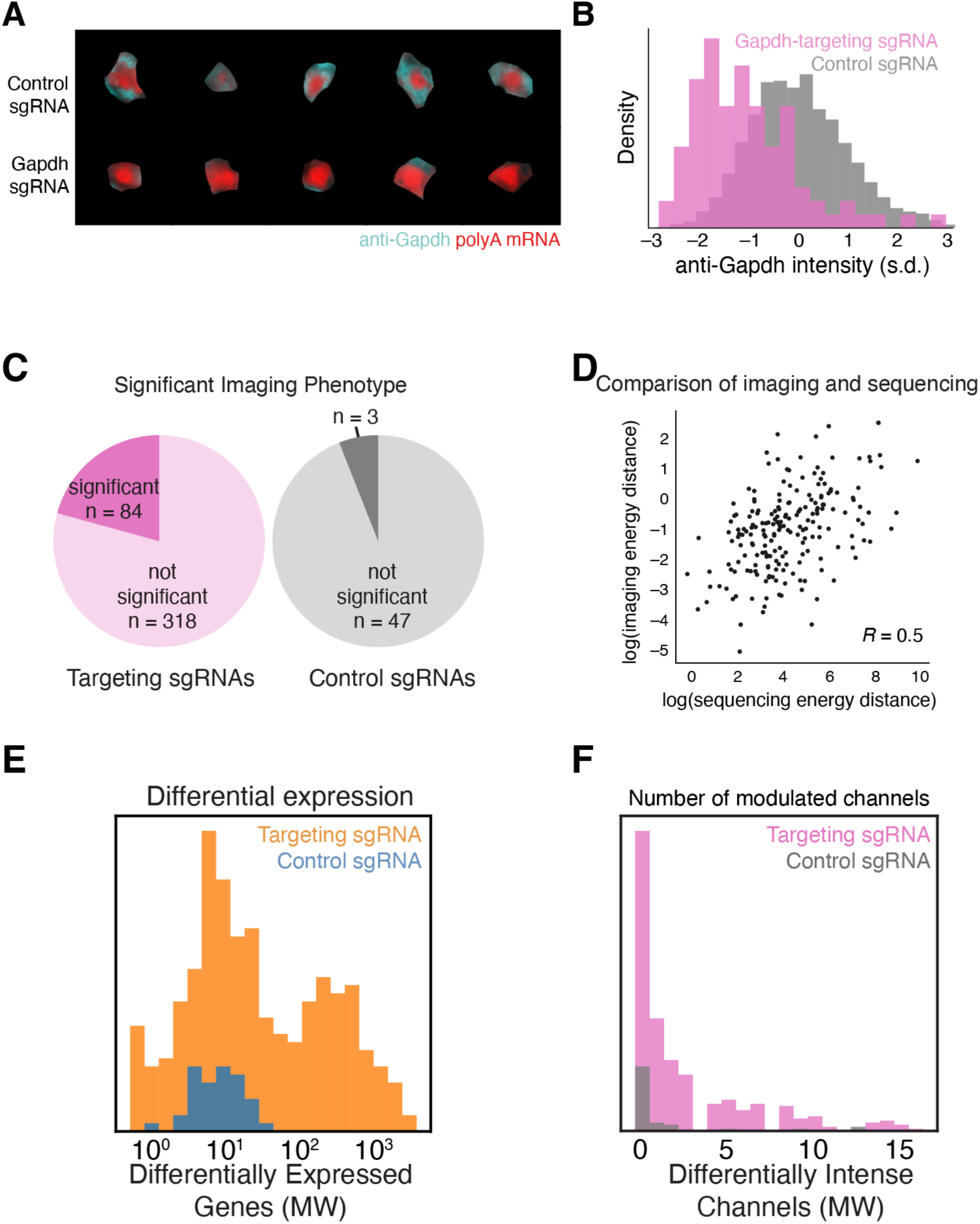
Additional analyses of perturbation data A. Unbiased sampling of cells with control sgRNAs and sgRNAs targeting *Gapdh*. The fluorescence micrographs show anti-GAPDH and polyA FISH channels. B. Histogram comparing anti-Gapdh intensity in called cells with a control sgRNA and called cells with *Gapdh*-targeting sgRNAs, from the imaging dataset. C. Pie charts showing the number of targeting (left) and control (right) sgRNAs that caused a significant transcriptional phenotype, as measured by a Holm-Sidak-corrected energy distance permutation test (p < 0.05), in the imaging dataset. D. Scatterplot comparing the energy distance vs control cells for each knockout in the imaging and Perturb-seq datasets. E. Histogram representing the number of differentially expressed genes for each perturbation, from the sequencing experiment. Differential gene expression reflects Benjamini-Hochberg-corrected, Mann-Whitney p < 0.05, versus cells with control sgRNAs. F. Histogram representing the number of imaging channels (proteins or RNAs) exhibiting differentially intense signals foreach perturbation, from the imaging experiment. Differentially intense signal reflects Benjamini-Hochberg-corrected, Mann-Whitney p < 0.05, versus cells with control sgRNAs.

